# Featural representation and internal noise around the visual field

**DOI:** 10.64898/2026.07.30.741910

**Authors:** Shutian Xue, Michael Landy, Marisa Carrasco

## Abstract

In human adults, visual performance varies systematically around the visual field. It is higher along the horizontal than the vertical meridian (horizontal–vertical anisotropy, HVA) and higher at the lower than the upper vertical meridian (vertical–meridian asymmetry, VMA). Although these robust performance fields have been linked to non-uniform neural resources, the system-level computations that translate neural constraints into perceptual asymmetries remain largely unexplored. Here, we used reverse correlation to characterize feature weighting and internal noise during peripheral orientation detection. Reverse correlation revealed non-ideal feature weighting in the joint orientation–spatial-frequency space, which was incorporated into a noisy-observer model jointly constrained by trial-wise detection responses and double-pass consistency. Across observers, the magnitude of the HVA in contrast sensitivity was correlated with individual asymmetries in orientation sensitivity and additive internal noise. In contrast, we found limited evidence that any tested representational or noise components reliably accounted for individual differences in VMA magnitude. Spatial-frequency tuning exhibited substantial individual variability, often peaking below the signal’s spatial frequency, but did not vary systematically across locations or explain performance asymmetries. These findings suggest that the HVA reflects systematic variation in the feature weighting of task-relevant orientations and internal noise, while constraining which computational components provide robust explanations of polar-angle asymmetries. Moreover, our framework provides a principled approach that links this prevalent perceptual asymmetry to the system-level computations that transform sensory information into perceptual decisions.

**Author summary:** Human vision is surprisingly uneven across our field of view. At the same distance from where we look, vision is better along the horizontal axis than the vertical axis, and better in the lower than the upper half of the vertical axis. These differences are well established, but little is known about their computational basis. To investigate, we combined visual detection tasks with computational modeling. By analyzing how observers detected faint patterns within noisy images, we measured how they process distinct features—like patterns with different orientations and spatial frequencies at different visual field locations. We then fit a mathematical model to separate distinct sources of internal noise. We found that differences in how the brain processes line orientations predicted the magnitude of the horizontal-vertical asymmetry across individuals, whereas distinct components of internal noise predicted these asymmetries in different ways. Our results show that visual field asymmetries are not driven by a single visual bottleneck, but rather by location-specific combinations of how the brain encodes relevant feature information and neural noise. This study helps explain why human vision is fundamentally uneven across our field of view.

## Introduction

Human visual performance declines with eccentricity [1, 2] and varies with polar angle, even at isoeccentric locations. Performance is better along the horizontal than the vertical meridian—termed the horizontal– vertical anisotropy (HVA)—and better along the lower than the upper vertical meridian—termed the vertical-meridian asymmetry (VMA) (Figure 1A; for a review, see [3]). In some tasks, the performance difference between the horizontal and vertical meridians can approach the magnitude of doubling or tripling eccentricity [4–6]. Both asymmetries are evident across fundamental dimensions, including contrast sensitivity [4–12] and spatial resolution/acuity [13–16].

**Fig. 1.**
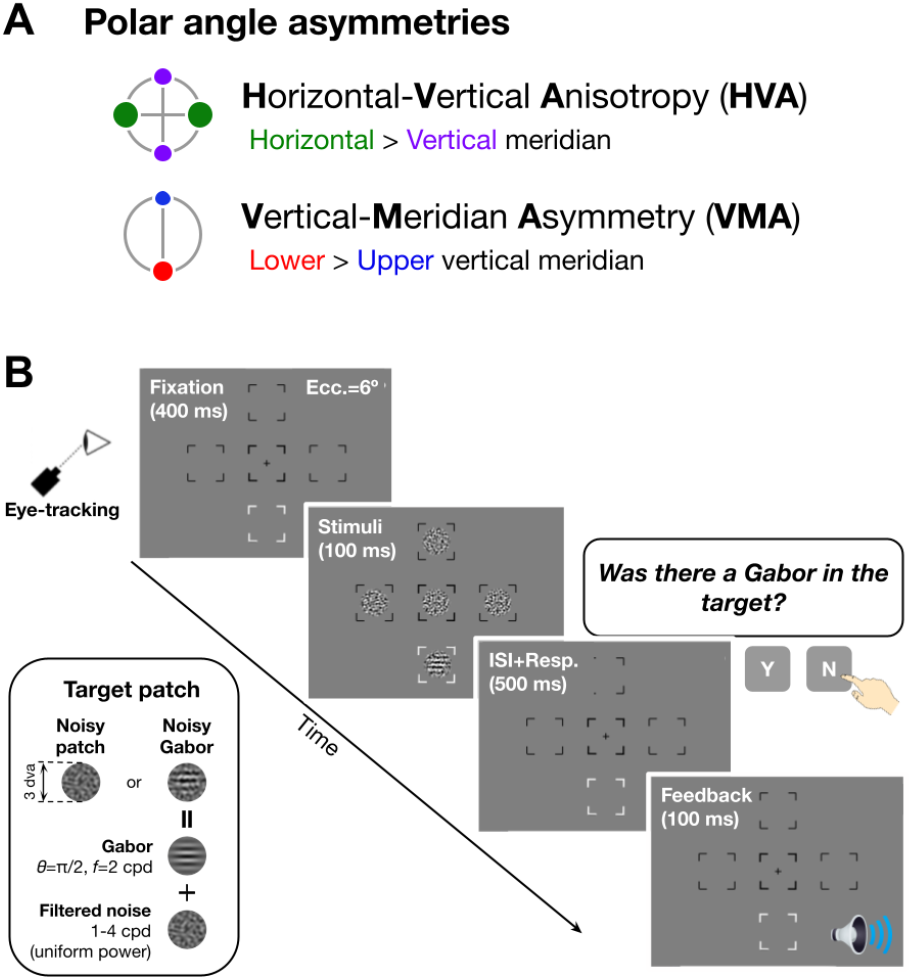
Polar angle asymmetries and trial sequence. **(A)** Performance varies around polar angle. It is better along the horizontal than the vertical meridian (HVA), and is better along the lower-than the upper-vertical meridian (VMA). **(B)** In a human behavioral experiment, each trial began with a central fixation cross and five placeholders displayed at the center and four isoeccentric locations along the cardinal meridians (6° eccentricity: left and right horizontal meridians, upper and lower vertical meridians). Subsequently, five stimulus patches, comprising one target and four distractors, were presented within the placeholders. On half of the trials, the target stimulus (indicated by a white placeholder) was a horizontal Gabor patch (2 cycles/deg or cpd, SD=0.8) embedded in filtered noise (spatial frequency: 1-4 cpd, RMS contrast: 20%); on the other half, the stimulus was an independently generated patch of filtered noise. The target location was constant within a block and varied across blocks. Observers reported whether the Gabor patch was present or absent by pressing one of two keys. Auditory feedback followed each response. The signal contrast was titrated to maintain approximately 70% accuracy.

### The limits of the “neural count” hypothesis

Understanding what gives rise to these perceptual patterns is crucial for revealing how the visual system operates under neural constraints and for building accurate models. A prominent “neural count” hypothesis posits better performance at locations with greater retinal and cortical representation, consistent with denser retinal sampling along the horizontal meridian [17–20] and larger primary visual cortex (V1) surface area that covaries with polar-angle performance [3, 21–25]. However, retinal and cortical “neural count” factors appear insufficient to fully account for polar-angle asymmetries. An image-computable model of orientation discrimination incorporating optical and retinal inhomogeneities explains only a limited fraction of behavioral variance—approximately 40% for the HVA and 10% for the VMA [26, 27]. Moreover, M-scaling stimulus size according to cortical magnification eliminated the eccentricity effect, but only reduced polar-angle asymmetries [6]. Together, these findings suggest that quantitative differences in retinal and cortical sampling alone do not fully explain polar-angle asymmetries, motivating the search for additional computational mechanisms.

Anatomical and physiological asymmetries do not specify how differences in neural resources are converted into behavioral performance. Greater neural representation could improve performance by increasing the amount of information ([28, 29]) or strengthening the population-level encoding of task-relevant stimulus features, which may in turn yield a stronger or more selective system-level perceptual template. Alternatively, it could reduce the internal noise that corrupts the decision variable, thereby increasing response consistency [30, 31]. These system-level computations are computationally distinct, but whether they play functionally distinct roles in shaping performance around polar angle is unknown. Dissociating them requires jointly characterizing how stimulus features are weighted and how internal noise limits the resulting decision process.

### System-level computations

This study tested whether polar-angle asymmetries reflect differences in system-level computations that transform noisy sensory input into perceptual decisions. We operationalized these computations as (i) the featural template used to represent task-relevant information, estimated via psychophysical reverse correlation, and (ii) internal noise that corrupts the resulting decision variable, constrained by trial-wise detection responses and double-pass response consistency in a noisy-observer model. Building on standard observer models of early vision [32–37], we asked whether polar-angle asymmetries arise from differences in the featural representation (i.e., the perceptual template), differences in internal noise, or both.

Reverse correlation captures how trial-to-trial fluctuations in stimulus features influence observers’ responses by leveraging the statistics of the visual input [38–41]. This method is an extension of the classification-image approach [36, 42, 43] with the addition of a generalized linear model (GLM; [37]). We focused on orientation and spatial frequency, two core features jointly encoded in early visual cortex (e.g., V1) (e.g., [44–47]). We summarize featural representation with tuning functions: higher gain indicates greater sensitivity to the task-relevant feature, whereas narrower bandwidth indicates greater selectivity. Tuning functions derived from reverse correlation have been shown to relate to sensory tuning properties measured in V1 [36], making reverse correlation a useful tool for linking behavior to feature-level encoding. Because featural representation differs across eccentricity [48], variation in featural representation around polar angle could contribute to the HVA and VMA. We therefore focused on whether polar-angle differences in sensitivity and selectivity predict polar-angle differences in contrast sensitivity.

The second aspect we examined—internal noise—is a fundamental constraint on sensory encoding and is typically reflected in reduced performance and reduced response reliability [30, 31, 49–52]. Although internal noise increases with eccentricity [48, 53, 54], whether it differs systematically around polar angle is unknown. We therefore leveraged a double-pass design to measure response consistency by presenting the same stimuli twice, which provides a direct behavioral constraint on internal noise [31, 55]. Within a noisy-observer model, we further distinguished noise that is private to each pass (“private noise”) from fluctuations shared across passes (“shared noise”), enabling us to assess their respective variation around polar angle and contribution to the HVA and VMA.

We first establish behavioral HVA and VMA in contrast sensitivity, then test whether these asymmetries are explained by polar-angle differences in orientation and spatial-frequency weighting, and finally determine whether decision-stage internal noise provides an additional constraint on performance. We show that, whereas behavioral HVA and VMA are robust, their computational signatures are distinct: HVA is linked to orientation sensitivity and additive internal noise, whereas VMA is not reliably explained by any of the representational or noise components tested here. Spatial-frequency weighting showed substantial individual variability but did not account for either asymmetry.

## Results

Observers performed a signal-detection task at isoeccentric locations along the horizontal meridian and the upper and lower vertical meridians (Figure 1B). Signal contrast was titrated to maintain stable accuracy across locations and was converted to contrast sensitivity to index performance. To dissociate the contributions of featural representation and internal noise to polar-angle asymmetries, we analyzed the behavioral data using two complementary approaches. First, we used psychophysical reverse correlation to estimate observer- and location-specific weighting over orientation and spatial frequency (Figure 2A) and tested whether asymmetries in these weights predicted individual differences in behavioral asymmetry. Second, we incorporated the resulting perceptual templates into a noisy-observer model to decompose internal noise into additive and shared components (Figure 2B). Finally, we evaluated which component of internal noise contributed to performance differences around polar angles. Specifically, we asked not only whether these computations differed systematically around polar angle, but also whether individual differences in their asymmetries predicted the magnitude of these performance asymmetries.

**Fig. 2.**
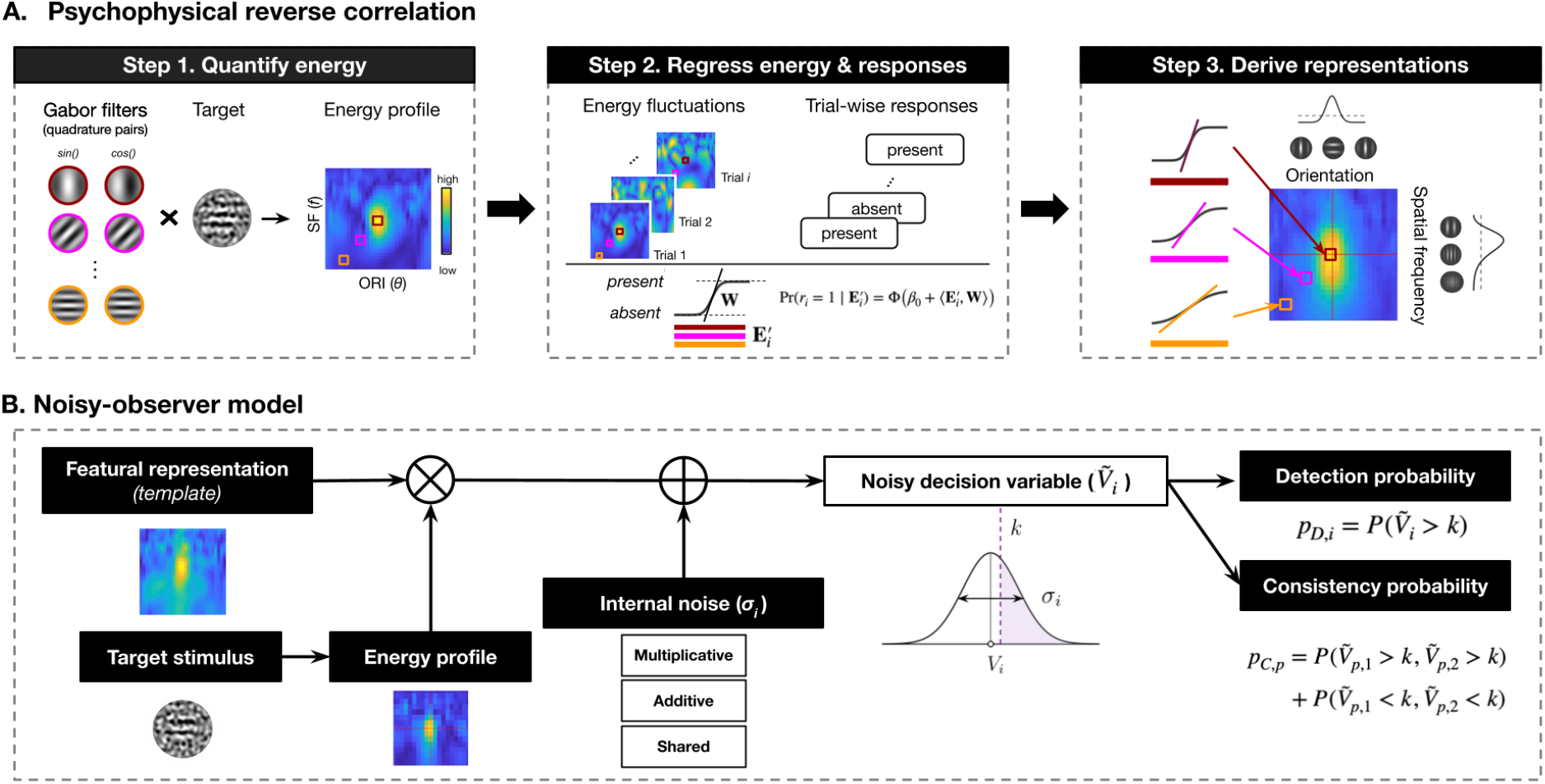
Reverse correlation and noisy-observer model schematic. **(A)** Reverse correlation was applied separately to the data from each observer and visual-field location. For each target stimulus, we computed an orientation *×* spatial-frequency contrast-energy profile by taking the inner product of the stimulus with a bank of quadrature-phase Gabor filter pairs (Step 1). For each feature combination, responses from the two filter phases were combined as root-sum-of-squares energy, yielding a two-dimensional energy profile for each trial. We then fitted a multivariate generalized linear model with a probit link, using the normalized trial-wise energy profiles to predict binary responses (“present” vs. “absent”) (Step 2). The resulting regression coefficients defined a two-dimensional perceptual weight map over orientation and spatial frequency. Finally, we marginalized the fitted two-dimensional weight map across each dimension to obtain one-dimensional orientation and spatial-frequency tuning functions (Step 3). Red vertical and horizontal lines indicate the signal orientation and spatial frequency, respectively. **(B)** Noisy-observer model. For each trial, the model computed a scalar decision variable as the inner product of the energy profile of the target stimulus and the observer-specific perceptual template estimated by reverse correlation. The decision variable was corrupted by additive and multiplicative noise private to each pass and by noise shared across the two presentations of a double-pass pair. The resulting noisy decision variable was compared with an observer- and location-specific decision criterion to generate a “present” or “absent” response. Additive-, multiplicative-, and shared-noise parameters and the criterion were estimated jointly from trial-wise detection responses and pairwise double-pass consistency.

### Contrast sensitivity shows HVA and VMA despite similar consistency and accuracy

Contrast sensitivity—the reciprocal of the signal contrast required to reach 70% accuracy—varied systematically with polar angle (Figure 3A). For planned comparisons, we averaged contrast sensitivity at the left and right locations to define the horizontal meridian, and averaged the upper and lower locations to define the vertical meridian. A one-way repeated-measures ANOVA across the horizontal meridian, lower-vertical, and upper-vertical meridians showed a location effect (*F* (2, 22) = 12.93, *p <* .001, 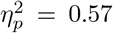, CI_95_ = [0.40, 0.76]). Planned pairwise comparisons showed that contrast sensitivity was higher at the horizontal than at the vertical meridian (*p <* .01, *g* = 0.86, CI_95_ = [0.40, 1.54]), indicating a horizontal–vertical anisotropy (HVA). Contrast sensitivity was also higher at the lower than at the upper vertical meridian (*p <* .001, *g* = 1.39, CI_95_ = [0.98, 2.35]), indicating a vertical-meridian asymmetry (VMA). These results confirmed well-established performance-field patterns in our data, setting the stage to link these performance variations to system-level computations.

**Fig. 3.**
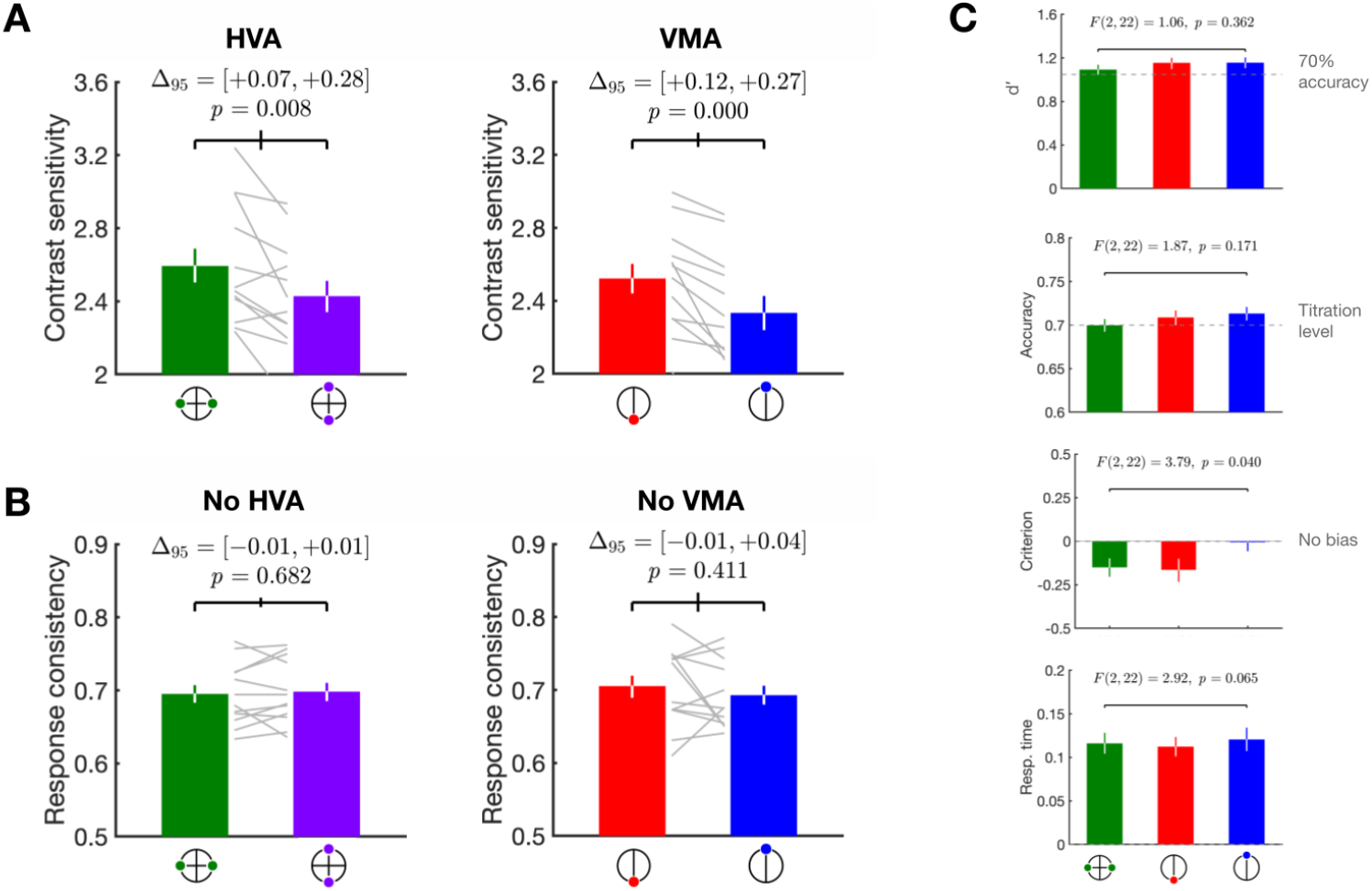
Behavioral metrics around polar angle. **(A)** Contrast sensitivity (the reciprocal of the contrast threshold yielding 70% accuracy) shows a robust horizontal–vertical anisotropy (HVA) and vertical-meridian asymmetry (VMA). Bars show the median of 10,000 bootstrapped group means (observers were resampled with replacement); error bars show 68% bootstrapped confidence intervals (CIs). For planned comparisons, metrics at the left and right locations were averaged to form the horizontal meridian, and metrics at the upper and lower locations were averaged to form the vertical meridian. Gray lines show individual observers (median across 1,000 cross-validated iterations). For each comparison, we report the between-location difference (median and 68% CI of the group-mean difference across observers). **(B)** Response consistencies (the proportion of identical responses across double-pass presentations) do not vary around polar angle. Plotting conventions are identical to panel A. **(C)** *d*^*′*^, accuracy, and response time are comparable around polar angle, indicating no location dependence in overall task difficulty or speed–accuracy trade-offs. Criterion is near zero at the upper vertical meridian and negative at other locations. Plotting conventions are identical to panel A.

We embedded a double-pass procedure in the main design to measure response consistency, as all trials were presented twice within each block. Response consistency—how often observers gave the same response to the repeated stimulus—did not differ around polar angle (both *p >* .1, Figure 3B), suggesting comparable overall internal noise across locations. In subsequent analyses, we used a modeling approach to decompose internal noise into distinct sources, with response consistency providing a key constraint on model fitting.

We verified that *d*^*′*^, accuracy, and response time were similar around polar-angle locations (Figure 3C), making it less likely that any location-dependent differences in overall task difficulty or speed–accuracy tradeoffs drove differences in featural representation. A one-way ANOVA showed that criterion had a location effect (*F* (2, 22) = 3.79, *p <* .05, 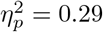, CI_95_ = [0.04, 0.66]): observers were nearly unbiased at the upper vertical meridian (criterion = −0.01, CI_95_ = [−0.11, 0.09]) and more biased toward responding that the signal was present at the horizontal meridian (criterion = −0.15, CI_95_ = [−0.25, −0.05]) and lower vertical meridian (criterion = −0.16, CI_95_ = [ 0.30, 0.04]). This criterion variation is unlikely to affect our main conclusions because our reverse-correlation analysis estimates the slope of the feature–response relation, whereas criterion primarily shifts the intercept (Figure 2A, *Step 2* ).

### Distinct featural representation varies around polar angle and contributes to performance asymmetries

We used psychophysical reverse correlation to estimate the featural representation at each polar-angle location, yielding a two-dimensional mapping of weights over orientation and spatial frequency (Figure 4A; see technical details in Methods: Reverse correlation). Warmer colors indicate larger positive weights, meaning that greater energy in a given orientation–spatial-frequency channel increased the likelihood that observers reported the signal as present. These weights, therefore, index the system’s sensitivity to information in each feature channel.

**Fig. 4.**
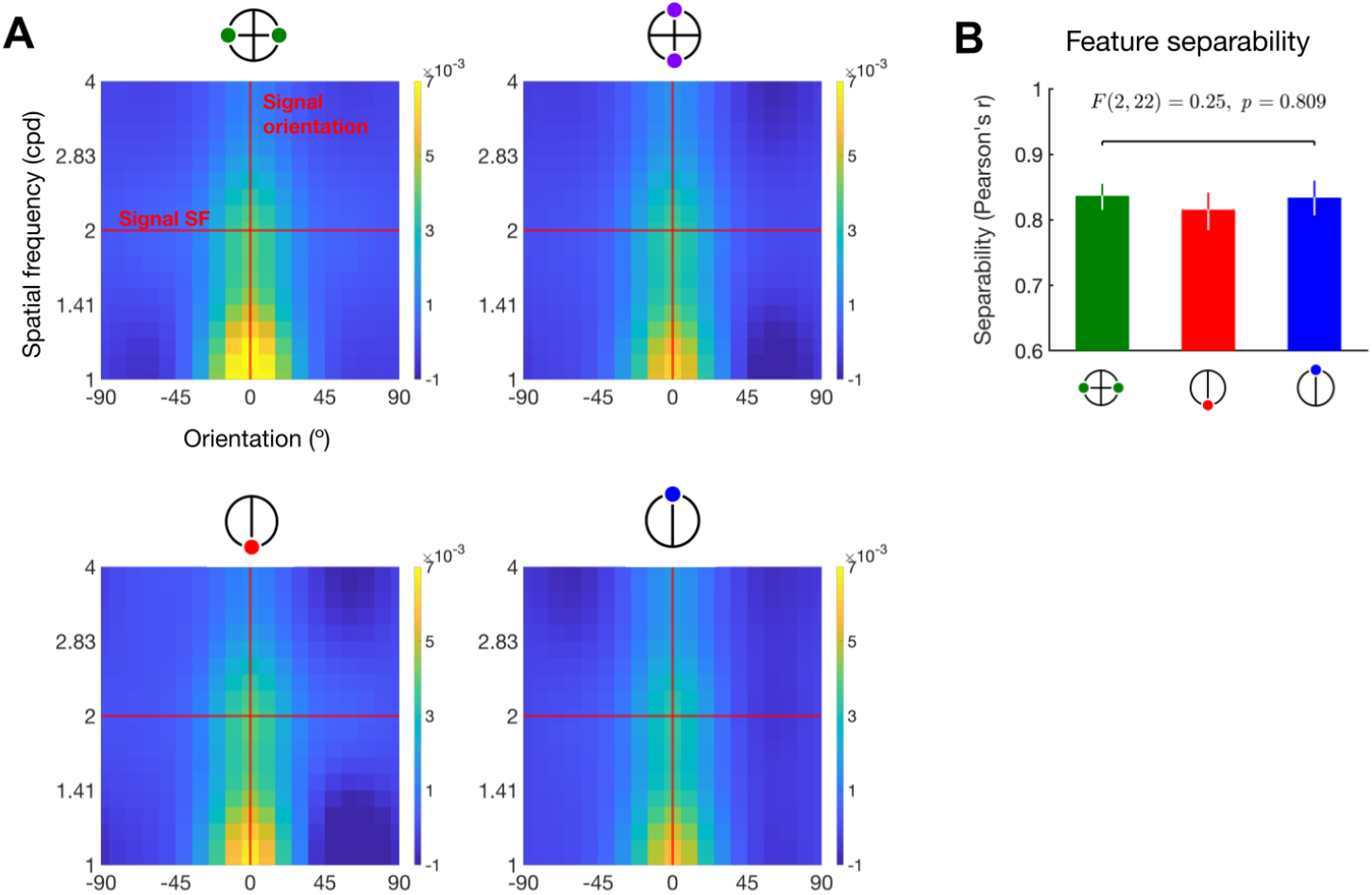
Featural representation around polar angle and feature separability. **(A)** Group-averaged two-dimensional featural representation over orientation and spatial frequency at each location. The map shown is the median of 1,000 cross-validated maps averaged across observers. Vertical and horizontal reference lines indicate the signal orientation and signal spatial frequency, respectively. For planned comparisons, representations at the left and right locations were averaged to form the horizontal meridian, and representations at the upper and lower locations were averaged to form the vertical meridian. **(B)** Feature separability. Plotting conventions are identical to Figure 3.

Across locations, the weight maps peaked near the signal orientation and at approximately 1 cycle per degree, below the signal spatial frequency of 2 cycles per degree. Our primary analyses focused on the height and bandwidth of the orientation-tuning peak. Qualitatively, weights appeared larger at the horizontal than at the vertical meridian and at the lower than at the upper vertical meridian, as reflected by the warmer colors in those maps. To quantify these differences separately for each feature dimension, we marginalized the two-dimensional representation over each axis to obtain one-dimensional orientation and spatial-frequency tuning functions. We quantified feature separability as the correlation between the two-dimensional featural representations and their rank-1 reconstruction (the outer product of the marginalized orientation and spatial-frequency tuning functions). High separability indicates that the marginalized tuning functions provide a faithful summary of the full two-dimensional mapping. We found that the two feature dimensions were highly separable (median *r >* 0.8; Figure 4B). Moreover, separability did not differ across locations (one-way repeated-measures ANOVA: *F* (2, 22) = 0.25, 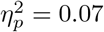, CI_95_ = [0.00, 0.32]). Thus, these marginalized tuning functions provide a faithful summary of the full 2D featural representation. We next quantified key properties of the tuning functions and compared them across paired locations.

#### Orientation tuning

We first characterized polar-angle differences in the orientation dimension. Figure 5 shows the marginalized orientation tuning functions averaged across observers, along with von Mises fits, for two comparisons: horizontal vs. vertical meridian (Figure 5A) and lower vs. upper vertical meridian (Figure 5F). From each fit, we compared the gain (height above baseline) and bandwidth (full-width at half-height) between paired locations (see Methods: Tuning function characterization). A larger gain indicates greater sensitivity, whereas a smaller bandwidth indicates sharper selectivity. At the group level, the horizontal and vertical meridians did not differ in gain (*p >* .1, *g* = 0.31, CI_95_ = [−0.08, +0.97]; Figure 5B) or bandwidth (*p >* .1, *g* = −0.09, CI_95_ = [−0.55, +0.65]; Figure 5C), indicating comparable sensitivity and selectivity. The lower and upper vertical meridians did not differ in gain (*p >* .1, *g* = 0.31, CI_95_ = [−0.26, +0.98]; Figure 5G) or bandwidth (*p >* .1, *g* = −0.11, CI_95_ = [− 0.87, +0.41]; Figure 5H). Together, these results indicate that at the group level, neither orientation sensitivity nor selectivity varied around polar angle.

**Fig. 5.**
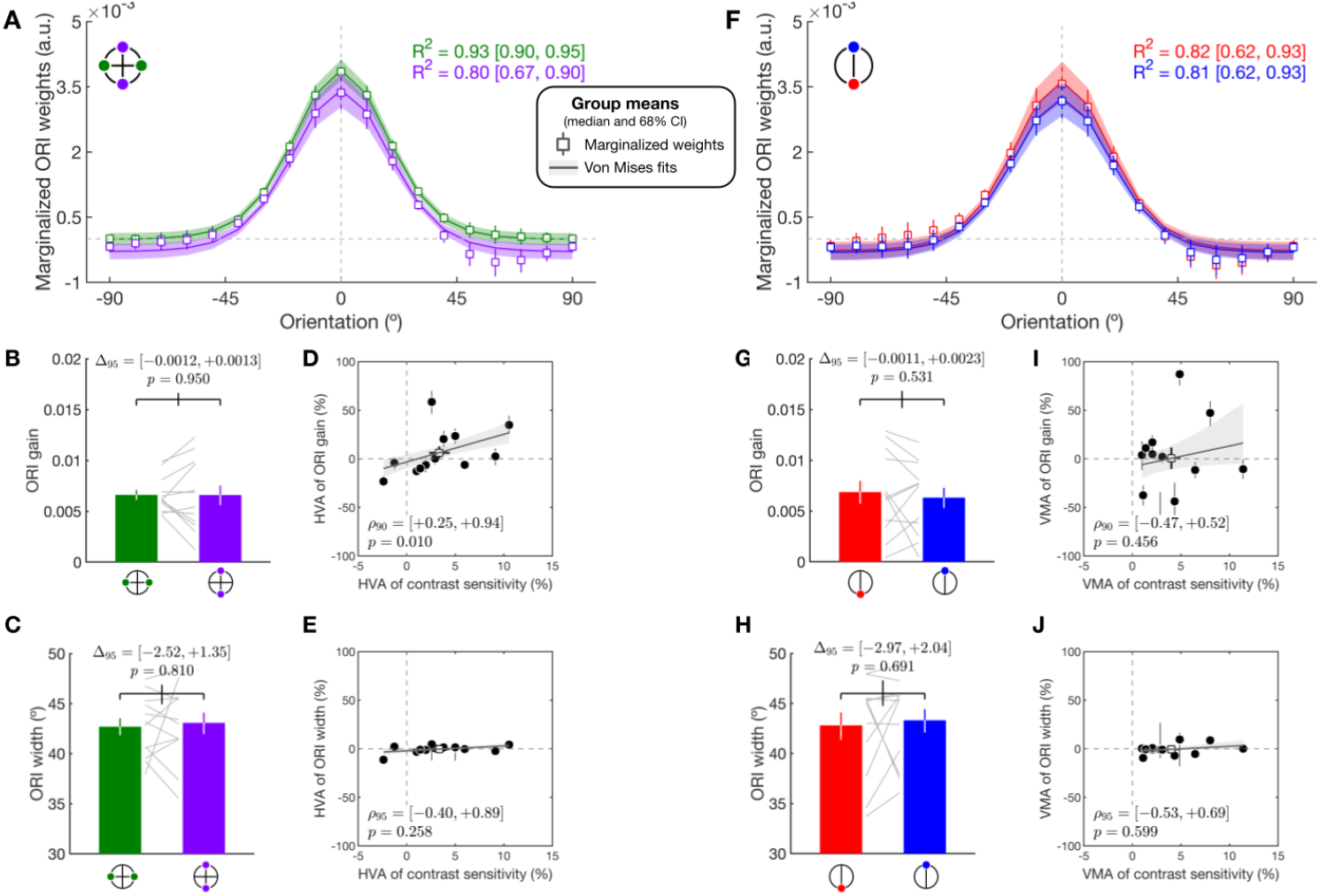
Orientation tuning functions and tuning parameters around polar angle. **(A)** Marginalized orientation tuning functions for the horizontal and the vertical meridian. 0° indicates the horizontal orientation. Squares and error bars show the median and 68% CI of bootstrapped group means, respectively. Curves and shaded bands show fits of a von Mises function (median and 68% CI of bootstrapped group fits). **(B–C)** Orientation tuning parameters derived from the fitted functions (gain and bandwidth). Plotting conventions are identical to Figure 3A. **(D–E)** Across observers, the association between HVA magnitude in orientation tuning parameters and HVA magnitude in contrast sensitivity. Dots and error bars show the median and 68% CI across cross-validated iterations for each observer. Solid lines and shaded bands show linear regression fits (median and 68% CI of bootstrapped fits). Dashed gray lines indicate zero difference. Spearman’s correlation coefficients are reported as the median and CI (CI range is defined a priori for correlation analyses) and permuted *p*-values are reported. **(F–H)** Same format as Panels A–E, but comparing the lower and upper vertical meridians to assess VMA-related tuning differences and their relationship to VMA magnitude in contrast sensitivity.

However, group-level location differences in tuning functions are not sufficient to establish a contribution to behavioral asymmetries. We therefore asked whether individual differences in tuning functions track individual differences in contrast-sensitivity asymmetries. For each metric, we computed a normalized asymmetry index, defined as the difference between two locations divided by their sum. This index quantified either the HVA (horizontal vs. vertical meridian) or the VMA (lower vs. upper vertical meridian). Our central question was whether asymmetry in gain or bandwidth predicted asymmetry in contrast sensitivity across observers.

To pre-specify the expected direction of the asymmetry correlations, we first quantified how each tuning parameter related to contrast sensitivity across observers while controlling for location (Figure 6). Contrast sensitivity was positively associated with orientation gain (*p <* .05; Figure 6A), indicating that better performance is linked to higher sensitivity to the preferred orientation. These modulation effects predict that asymmetry in contrast sensitivity should increase with asymmetry in gain, providing an a priori basis for one-tailed tests in the asymmetry correlations below. Consistent with a one-tailed test at *α* = 0.05, we report two-tailed 90% confidence intervals for these directional analyses. For the remaining non-directional correlation analyses, we report standard two-tailed 95% confidence intervals.

**Fig. 6.**
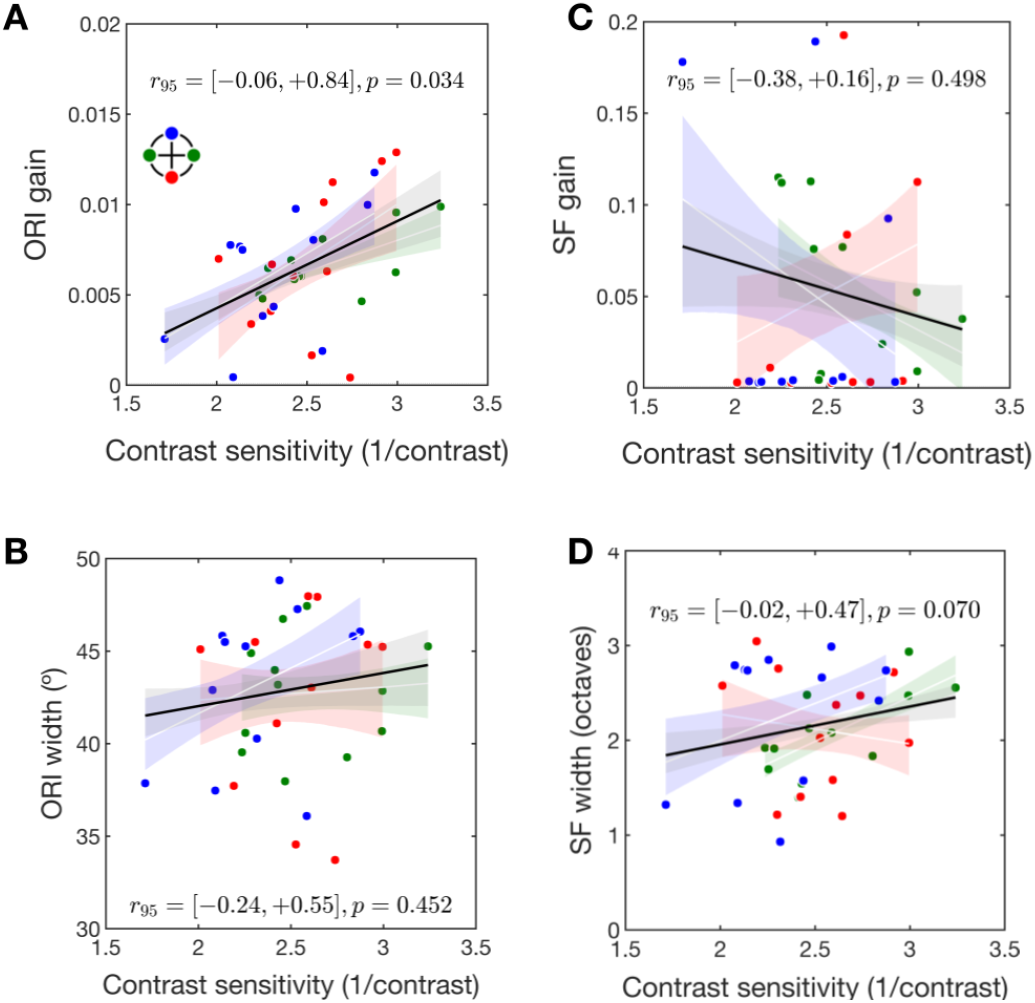
Correlations between contrast sensitivity and tuning parameters across observers and locations. **(A)** Scatterplot of observers’ orientation gain plotted against contrast sensitivity across the horizontal, lower vertical, and upper vertical meridians. Dots show the median across cross-validated iterations for each observer. Solid lines and shaded gray bands show linear regression fits across all locations (median and 68% CI of bootstrapped fits). Colored shaded bands show within-location regression fits (68% CI of bootstrapped fits). The partial correlation coefficient (median and 95% CI) is reported at the bottom. **(B–D)** Same format as Panel A for orientation tuning bandwidth, spatial-frequency gain, and spatial-frequency tuning bandwidth, reporting the partial correlation coefficient with its corresponding 95% CI.

We observed a dissociable pattern of associations between the computations underlying the HVA and the VMA. For the HVA, asymmetry in contrast sensitivity was positively associated with asymmetry in gain (*p <* .05; Figure 5D), but not with asymmetry in bandwidth (*p >* .1; Figure 5E). For the VMA, asymmetry in contrast sensitivity was not associated with asymmetry in gain (*p >* .1, Figure 5I) or bandwidth (*p >* .1; Figure 5J). Thus, in the orientation domain, HVA magnitude—but not VMA magnitude—tracked sensitivity differences.

Importantly, group-level mean differences and across-observer covariation address distinct questions and need not align. For example, orientation gain did not differ between the horizontal and vertical meridians at the group level (Figure 5B, also shown as black-edge squares in Figure 5D), yet individual differences in gain asymmetry predicted asymmetry in contrast sensitivity for the HVA (Figure 5D). This across-observer covariation provides a more specific test of whether a representational parameter explains a behavioral asymmetry.

#### Spatial-frequency tuning

We next applied the same analytic framework to the representation of spatial frequency to test whether polar-angle asymmetries also arise from differences in spatial-frequency sensitivity or selectivity. Figure 7 shows the marginalized spatial-frequency tuning functions, log-Gaussian fits, and the derived gain and bandwidth parameters. At the group level, neither gain (*p >* .1, *g* = +0.00, CI_95_ = [−0.73, +0.52]; Figure 7B) nor bandwidth (*p >* .1, *g* = −0.12, CI_95_ = [− 0.69, 0.49]; Figure 7C) differed between the horizontal and vertical meridians. Similarly, neither gain nor bandwidth differed between the upper and lower vertical meridians (*p*s *>* .1, *g <* 0; Figure 7G,H). Thus, at the group level, spatial-frequency tuning parameters did not differ around polar angle.

**Fig. 7.**
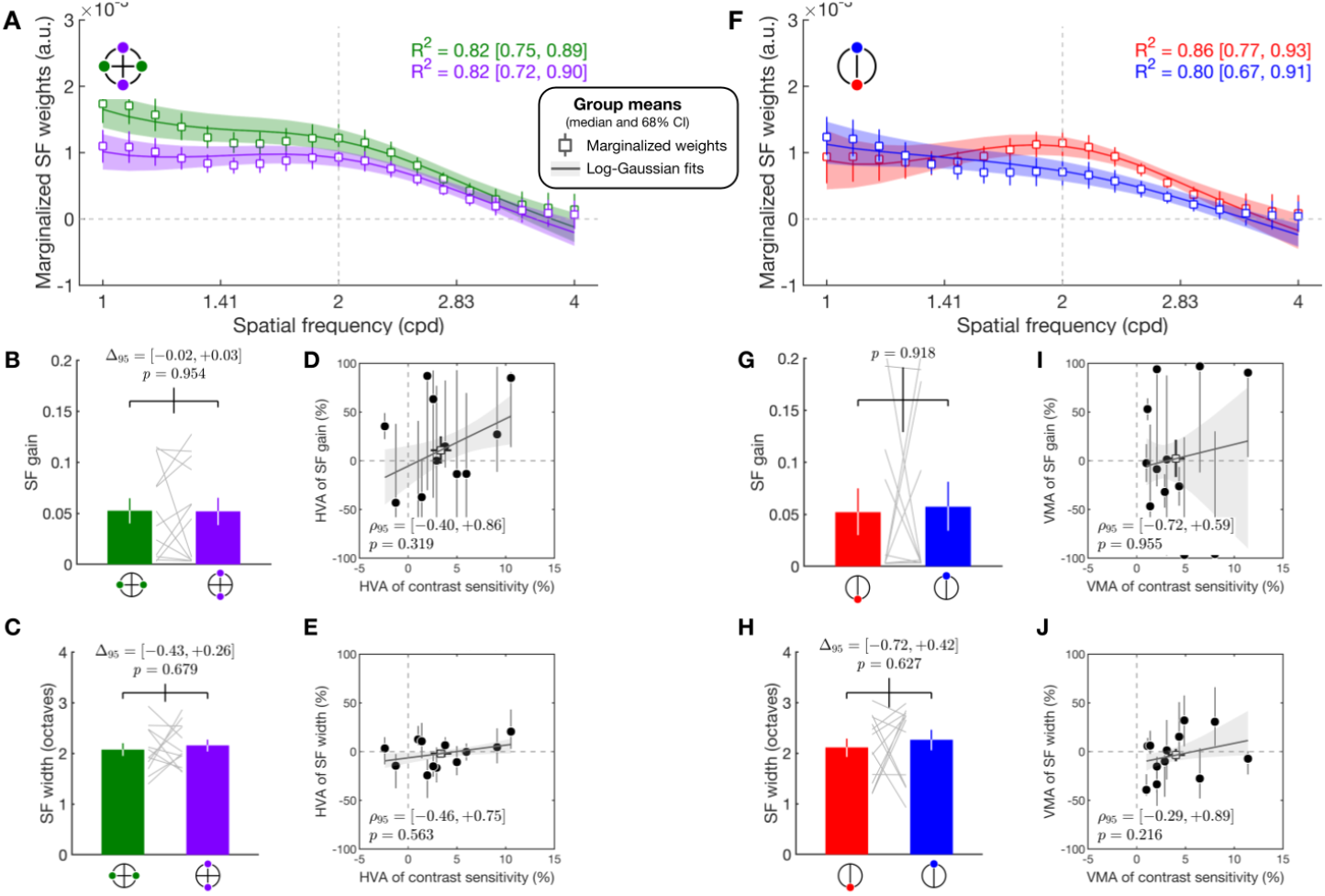
Spatial-frequency tuning functions and tuning parameters around polar angle. **(A)** Marginalized spatial-frequency tuning functions for the horizontal and vertical meridians. Squares and error bars show the median and 68% CI of bootstrapped group means, respectively. Curves and shaded bands show fits of a log-Gaussian function (median and 68% CI of bootstrapped group fits). (B–C) Spatial-frequency tuning parameters derived from the fitted functions (gain and tuning bandwidth). Plotting conventions are identical to Figure 3A. (D–E) Across observers, the association between HVA magnitude in spatial-frequency tuning parameters and HVA magnitude in contrast sensitivity. Plotting conventions are identical to Figures 5D–E. **(F–J)** Same format as Panels A–E, but comparing the lower and upper vertical meridians to assess VMA-related tuning differences and their relationship to VMA magnitude in contrast sensitivity.

As in the orientation analysis, we next tested whether individual differences in tuning parameters tracked individual differences in performance asymmetries. First, contrast sensitivity did not correlate with either spatial-frequency gain or bandwidth (Figure 6C,D). Because this analysis did not establish a clear directional prediction, we used two-tailed tests for the asymmetry correlations and reported 95% confidence intervals for the correlation coefficients. We found that asymmetry in contrast sensitivity did not track asymmetry in spatial-frequency gain or bandwidth for either the HVA or the VMA (Figure 7D,E,I,J). Thus, these spatial-frequency tuning parameters did not reliably explain individual differences in the HVA or VMA in this task.

### Noisy-observer model captures detection and consistency and yields behaviorally meaningful estimates

Having characterized how featural representation contributed to polar-angle asymmetries, we next estimated internal noise using a noisy-observer model that links each trial’s decision variable to both detection rates and response consistencies (Figure 8A; see Methods: Noisy-observer model). Empirical detection rates and response consistency are plotted as a function of the binned decision variable, along with predictions of the model that best captured the data (including additive and shared noise), computed by averaging trial-wise predicted probabilities within each bin. Across observers, detection rates increased monotonically with the decision variable (Figure 8A, left), consistent with larger decision-variable values reflecting more task-relevant information in the stimulus.

**Fig. 8.**
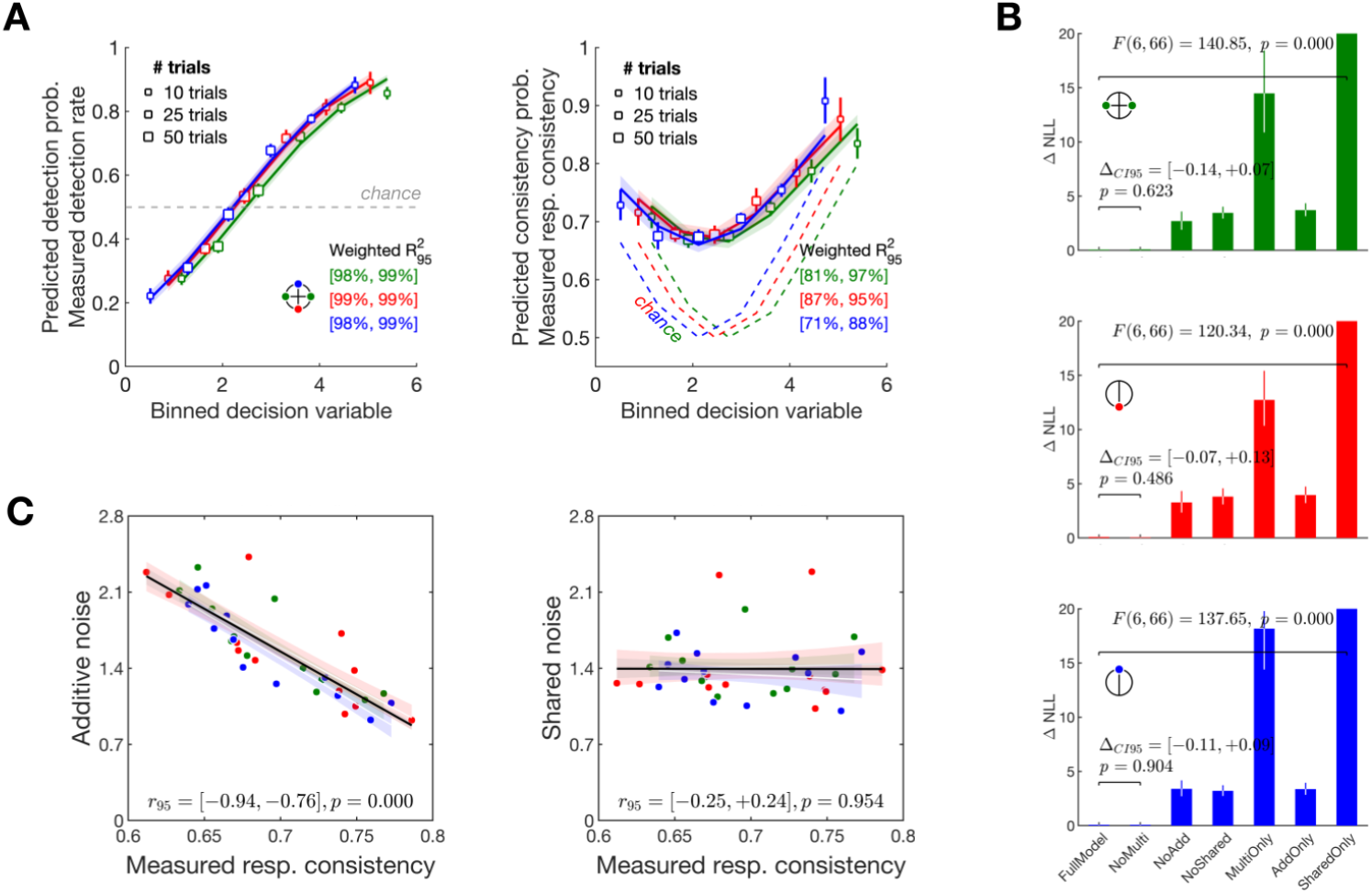
Noisy-observer model fits for detection, double-pass consistency, and behavioral relevance. **(A)** Empirical detection rate (left) and response consistency (right) as a function of the binned decision variable. Squares and error bars show the median and 68% CI of bootstrapped group means. Symbol size indicates the number of trials within each bin. Curves and shaded bands show predictions of the model that omits multiplicative noise (median and 68% CI of bootstrapped group predictions). *R*^2^ (median and 95% CI) is reported for each location. In the left panel, the dashed gray line marks the chance detection rate, *p*_*D,i*_ = 0.5. In the right panel, the colored dashed lines mark the expected response consistency assuming independent responses, 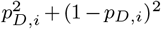. **(B)** Model comparison across locations. Model variants either include all parameters (*FullModel* ) or selectively exclude multiplicative (*NoMulti* ), additive (*NoAdd* ), or shared (*NoShared* ) internal noise, or only include one noise parameter (*MultiOnly, AddOnly, SharedOnly* ). Difference in negative log-likelihood (Δ NLL) is computed as each reduced model’s cross-validated negative log-likelihood minus the lowest value among all models at that location; lower values indicate better fits. Plotting conventions are identical to Figure 3A. **(C)** Scatterplot of observers’ internal noise plotted against response consistency across observers and locations. The partial correlation coefficient is reported within the panel. Plotting conventions are identical to Figure 6.

Response consistency showed a characteristic non-monotonic profile (Figure 8A, right): it was high when the decision variable was very low (clearly noise-only stimuli) or very high (strong signal-like structure), and lowest at intermediate values where observers were most uncertain. Critically, at intermediate decision-variable values, empirical consistency exceeded the independence baseline (dashed line), which shows the expected consistency under independent responses at the chance detection rate 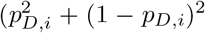 when *p*_*D,i*_ = 0.5). This excess consistency indicates trial-to-trial covariation between the two passes, which is accounted for by the shared-noise component in the model.

The model chosen to fit the data includes additive and shared noise but no multiplicative noise. For each observer and location, we computed ΔNLL as each model’s negative log-likelihood (NLL) minus the lowest NLL among all seven model variants at that location for that observer (Figure 8B). Thus, a lower ΔNLL indicates a better fit. A two-way repeated-measures ANOVA (model variant × location) revealed an interaction between the model variant and location (*F* (6, 66) = 152.45, *p <* .001, 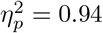, CI_95_ = [0.91, 0.97]). Planned within-location comparisons showed that the full model and the model without multiplicative noise outperformed all other variants for all locations (Figure 8B). Because these two models did not differ significantly from each other, we selected the simpler model without multiplicative noise. Predictions from this variant are shown in Figure 8A. Simulation-based recovery analyses further supported this choice: the reduced model without multiplicative noise showed good parameter recovery, whereas models containing both additive and multiplicative noise exhibited a tradeoff between these parameters (Appendix C).

Finally, we examined whether the estimated noise parameters tracked the response consistency. Figure 8C shows that across observers (controlling for location), response consistency decreased with higher additive noise (*p <* .001) and did not vary as a function of shared noise (*p >* .1). Thus, additive noise best captured behaviorally relevant response variability.

Together, these analyses show that the noisy-observer model jointly accounts for detection and double-pass consistency, and yields separable internal-noise components that capture response dependence across repeated presentations.

### Internal noise is spatially uniform but tracks HVA

We next asked whether model-derived internal noise parameters differ around polar angle. Neither additive nor shared noise differed between the horizontal and vertical meridians (*p >* .1, Figure 9A) or along the vertical meridian (*p >* .1, Figure 9B), in agreement with the constant response consistencies shown in Figure 3B.

**Fig. 9.**
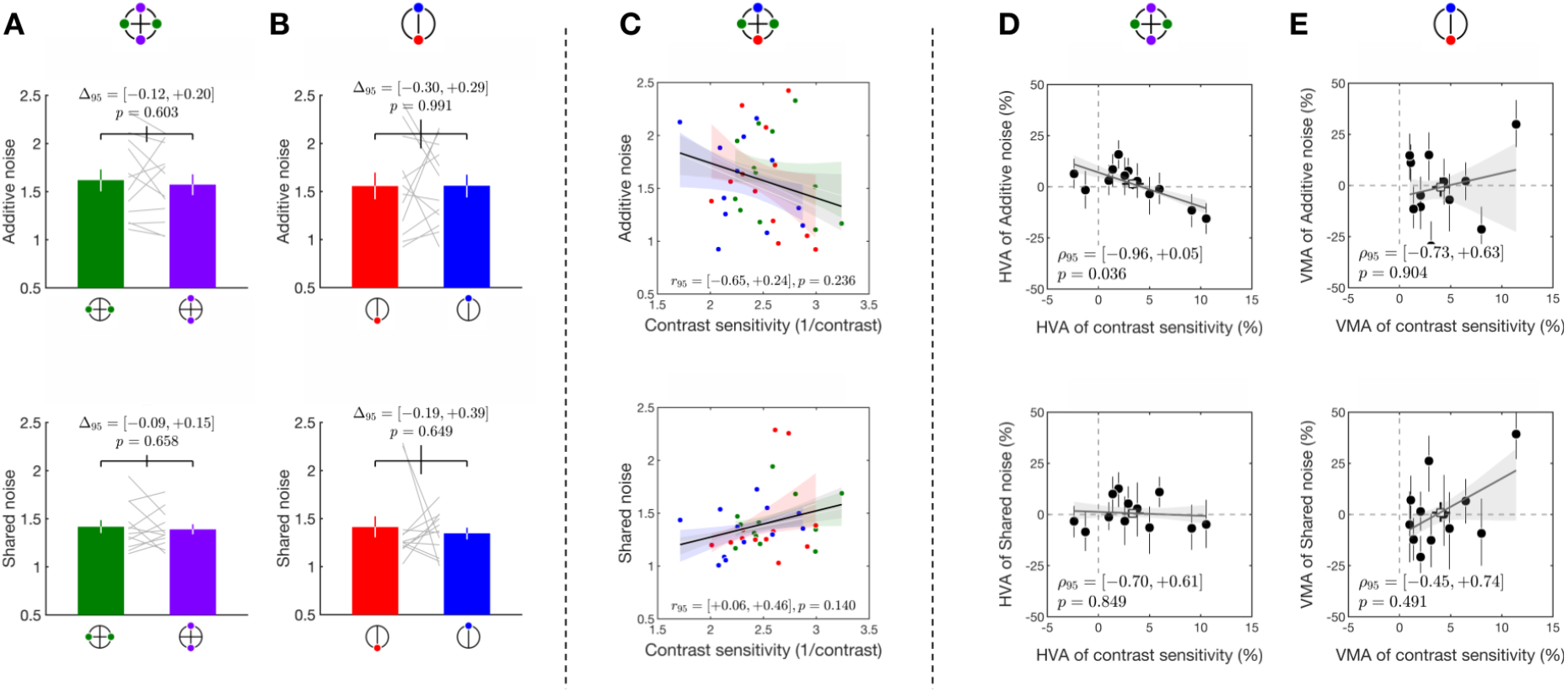
Internal noise components around polar angle and their behavioral relevance. **(A)** Estimated internal noise components at the horizontal meridian compared with the vertical meridian. Plotting conventions are identical to Figure 3A. **(B)** Same format as Panel A, but comparing the lower and upper vertical meridians. **(C)** Scatterplot of observers’ internal noise plotted against contrast sensitivity across locations. Plotting conventions are identical to Figure 6. **(D)** Across observers, the association between HVA magnitude in each internal noise component and HVA magnitude in contrast sensitivity. Plotting conventions are identical to Figure 5D–E. **(E)** Same format as Panel D for the VMA comparison.

We then asked whether internal-noise estimates related to contrast sensitivity (Figure 9C). Controlling for location, contrast sensitivity did not correlate with the amount of additive noise (partial Pearson’s *r* = −0.30, CI_95_ = [−0.65, +0.24], *p >* .1) or shared noise (partial Pearson’s *r* = +0.26, CI_95_ = [+0.06, +0.46], *p >* .1). Because these analyses did not establish a directional prediction, we tested asymmetries in both additive and shared noise using two-tailed correlations. Asymmetry in additive internal noise predicted the magnitude of the HVA (*p <* .01; Figure 9D), but not the VMA (*p >* .1; Figure 9E). This means that the magnitude of the HVA for contrast sensitivity—how much better performance was at the horizontal than at the vertical meridian—was predicted by the asymmetry of additive noise across those same meridians. In contrast, neither additive-noise asymmetry nor shared-noise asymmetry predicted VMA magnitude (Figure 9E). Together, these results indicate that although additive internal noise did not differ significantly between the horizontal and vertical meridians at the group level, individual variation in this asymmetry nevertheless tracked HVA magnitude in contrast sensitivity. Shared internal noise, in contrast, did not reliably track either performance asymmetry.

In summary, the HVA was associated with individual differences in both orientation sensitivity and additive internal noise: observers with stronger horizontal-meridian advantages in these measures showed larger HVA in contrast sensitivity. In contrast, spatial-frequency tuning did not reliably predict either the HVA or the VMA. None of the tested featural or noise measures reliably accounted for the VMA.

## Discussion

Peripheral detection was not governed by an ideal, signal-matched template: observers relied on a non-ideal feature-weighting readout strategy emphasizing spatial frequencies lower than those in the target, and their readout was limited by internal noise. Using reverse correlation and a noisy-observer model constrained by both detection performance and double-pass consistency, we tested whether mismatch of the template with the target, along with internal noise, explained polar-angle asymmetries in contrast sensitivity. The magnitude of the HVA was associated with individual asymmetries in orientation sensitivity and additive internal noise, whereas none of the tested components explained the magnitude of the VMA. Spatial-frequency tuning did not vary systematically across locations or account for either asymmetry. Together, these results suggest that the HVA, but not the VMA, reflects variations in featural representation and internal noise.

### Behavioral implications of feature weighting and internal noise

#### Noiseless orientation-related tasks

Our results offer a computational interpretation of the long-standing observation that performance fields around the polar angle are robust across tasks in which performance strongly depends on orientation-related structure, including detection of oriented patterns [4, 8] and categorical orientation discrimination [4–7, 9– 12, 15, 56, 57]. For noiseless stimuli, performance depends primarily on how strongly the orientation structure in the stimulus is weighted and how sharply it is represented. Locations with higher orientation sensitivity can generate a stronger template-matched decision variable for the same stimulus input, which increases the probability of a signal-consistent readout and improves accuracy. These computational differences naturally translate into behavioral findings such as higher accuracy [4, 57], higher inferred contrast sensitivity [5–11], and higher acuity [15, 16].

#### HVA and VMA should be treated as partially separable

Our results support a dissociable view of the HVA and VMA, which are sometimes discussed under a single “performance field” framework. Across observers, HVA magnitude covaried with asymmetries in orientation sensitivity and additive internal noise, whereas we found no compelling account for the VMA. These results suggest that the two asymmetries can reflect distinct components of the feature-weighting readout and internal noise inferred from behavior. This dissociation aligns with behavioral and neural evidence that the HVA and VMA are only weakly coupled: their magnitudes are often uncorrelated across individuals for contrast sensitivity [5, 10–12, 57] and acuity [15]. Moreover, individual HVA and VMA magnitudes are only marginally correlated with the V1 area devoted to each region of the visual field [24].

Developmental findings provide converging evidence that the HVA is present from childhood to adulthood in both behavior and cortical surface area, whereas the VMA is reduced or absent in children [58, 59]. Furthermore, the VMA develops gradually throughout adolescence, becoming as pronounced as in adults only during late adolescence [60]. This trajectory is consistent with the possibility that the computations supporting the VMA mature later and may be more experience-dependent. This idea is supported by anatomical evidence showing that cone density exhibits an HVA, but no VMA [19, 61]. Furthermore, the HVA in midget retinal ganglion cell (mRGC) density [18, 20], V1 surface area [22–24, 59], and perceptual performance [10, 15, 57] are more pronounced than the corresponding VMA.

#### Reconciling reverse-correlation and equivalent-noise approaches

Our results complement equivalent-noise and perceptual-template-model findings suggesting that polar-angle asymmetries primarily reflect differences in signal amplification [62]. First, whereas the perceptual-template model assumes a signal-identical ideal template, reverse correlation provides feature-level resolution by estimating how observers weight orientation and spatial-frequency information. From this perspective, location-dependent gain changes [62] may reflect location-dependent reweighting at the template-matching stage (current study). Thus, despite differences in task, stimulus statistics, and model structure, both approaches point to a shared bottleneck in how task-relevant evidence is filtered and matched to a decision template.

Second, the two approaches decompose internal noise differently: The perceptual template model infers noise from threshold changes across multiple external-noise levels, whereas the current model estimates decision-variable noise at a fixed external-noise level and includes a shared component constrained by double-pass consistency. Additive noise showed broadly consistent spatial patterns across studies, whereas neither framework provided strong evidence for multiplicative noise. This does not imply that multiplicative noise is absent; because signal strength was titrated in the present task, the decision-variable range was restricted, meaning that decision-variable-dependent noise may have had a limited dynamic range, allowing our results to be modeled with additive noise alone.

### System-level computations in feature weighting and internal noise

#### HVA is sensitivity-linked

One plausible account is that the HVA is especially sensitivity-linked because the horizontal meridian has stronger neural-resource advantages available to the readout. V1 surface area (and related cortical magnification) is larger along the horizontal than along the vertical meridian, and this horizontal-vertical difference is more pronounced than the difference between the upper and lower vertical meridians [22, 23, 59]. Individual differences in local V1 magnification correlate with perceptual sensitivity and acuity across observers [6, 11, 24, 63–65]. Under standard population-coding assumptions, engaging a larger cortical population can increase the effective signal-to-noise ratio of template-matched evidence, depending on the structure of correlated noise [28, 66, 67]. This account provides a plausible basis for a sensitivity-linked HVA, as differences in surface area could translate into differences in the sensitivity to task-relevant features (i.e., orientation) in the readout.

#### A more limited account of VMA

In contrast, our results did not provide compelling evidence for a mechanism underlying the VMA. One possibility is that VMA-related differences may depend more on feature selectivity than on feature sensitivity, but the present detection task may not have been optimized to reveal such effects. Population receptive field (pRF) sizes in early visual cortex vary around polar angle in a pattern that mirrors both the HVA and VMA [21, 59]. Smaller pRFs imply reduced spatial pooling and greater spatial precision, which could in principle support sharper readout of task-relevant features and manifest as higher selectivity in an inferred template.

However, because the current task required detecting the presence of oriented structure, and spatial-frequency information may have been less diagnostic under bandpass-filtered noise, performance may have depended more strongly on sensitivity to task-relevant evidence than on fine-grained feature selectivity. Thus, the absence of a reliable VMA-linked mechanism in the present data should not be taken as evidence that the VMA lacks a computational basis, but rather as a constraint on what this particular task and model could reveal.

### Interpretation of additive and shared noise

Private noise components—capturing stimulus-independent (additive) and stimulus-dependent (multiplicative) noise—are typically considered performance-limiting because they reduce the reliability of sensory evidence and thus effective decoding precision [28, 51, 68–70]. In contrast, shared noise—a less studied component—was found to be uniform around polar angle and did not contribute to performance asymmetries. One possible interpretation is that the shared term partly reflects a mismatch between the observer’s true internal template and the template recovered by reverse correlation. Such a mismatch could arise from residual channel correlations in the filtered-noise stimulus ensemble, trial-to-trial variation in the mechanisms that guide responses, or nonlinear computations not captured by the current model [71]. Because any mismatch would be relatively stable within an observer and location, it would affect both passes of a double-pass pair similarly and therefore appear as a shared source of noise. Importantly, this interpretation does not undermine the utility of the recovered template, which still provides a useful estimate of the observer’s effective feature weighting during task performance.

Because only the target stimuli were repeated across passes, we assume that signal-like energy fluctuations in the independently generated distractors did not systematically influence responses to the target location. Any such influence would instead contribute to reduced cross-pass consistency.

These noise terms should be interpreted as effective decision-stage quantities, not direct measurements of neural noise. They may aggregate multiple sources, including early sensory noise, cortical spiking variability, and fluctuations in brain state [51, 68, 69, 72–78]. Importantly, the shared-noise term should not be equated with correlated neural noise or spike-count correlations [28, 51, 68–70]. Here, it simply captures cross-pass dependence at the decision stage and may partly reflect stable template mismatch or nonlinear computations not captured by the model. Resolving how these effective noise components map onto neural mechanisms will require an explicit bridging model and matched neural measurements.

### Implications for neural constraints

A large body of neurophysiology and neuroimaging literature shows that early visual pathways are spatially non-uniform around polar angle, including retinal and LGN sampling [26, 27, 79], V1 cortical magnification and surface area [21–24, 27, 59], population receptive field (pRF) size [21, 59], and stimulus-evoked signals in early visual cortex (e.g., BOLD amplitude; [56, 80]). These anatomical and voxel-level measurements establish the existence of non-uniform neural resources, but they do not specify how that information is filtered and pooled to drive a perceptual decision. In principle, the sampling asymmetries described above could influence behavior through one or both of two routes: improving the quality of task-relevant evidence (i.e., yielding stronger or sharper template-matched signals) or reducing the effective internal noise that limits reliability. Our results help fill this missing link by providing a behavioral–computational characterization of the readout, with corresponding neural implications.

First, our results support the hypothesis that polar-angle performance fields are determined not only by how much cortex represents a location, but also by how that information is filtered and pooled into a decision variable. This framing matches the interpretation that differences in neural sampling provide an important substrate and can reduce asymmetries when matched via M-scaling, but they do not fully account for behavioral performance fields—implying the existence of additional readout constraints beyond non-uniform sampling [6, 26, 27]. Second, the effective feature weights inform the decision. Although these weights can resemble properties of single-neuron receptive fields (Neri & Levi, 2006), they are not meant to localize tuning at the single-neuron level; instead, they summarize population-level filtering and pooling in the task-relevant feature space, and are intended to guide mechanistic hypotheses that link neural population activity to behavior.

### Model novelty and limitations

Many models in early vision link a perceptual template to internal noise to explain detection and discrimination performance (e.g., [32–35]). Our approach extends this tradition in two ways. First, instead of assuming an ideal template, we estimate each observer’s effective template using reverse correlation and embed it directly in the noisy-observer model. Second, we constrain internal-noise estimates using both trial-wise detection responses and double-pass consistency. Together, these features allow us to dissociate the contributions of feature weighting and internal noise to visual-field asymmetries.

Two limitations define the scope of our conclusions. First, the estimated template and noise parameters are effective behavioral quantities and do not uniquely identify their underlying neural sources. Second, the template mismatch we measured likely reflects the well-documented reduction in sensitivity to higher spatial frequencies in the periphery [6, 39, 40, 62, 81, 82]. Establishing a mechanistic link between non-uniform neural sampling, feature weighting, and internal noise will require an explicit bridging model beyond behavior alone.

### Conclusion and implications

This study provides evidence that visual performance fields reflect systematic differences in feature representation and internal noise, rather than a uniform rescaling of a single computation across the visual field. By combining reverse correlation with a noisy-observer model, we characterized observer-specific feature weighting and effective decision-stage noise within a common framework. Our results indicate that individual differences in the HVA are associated with differences in orientation-based feature weighting and internal noise, whereas spatial-frequency weighting exhibited substantial variability across observers and did not reliably account for performance asymmetries.

More broadly, these findings suggest that visual performance fields arise from interactions between local neural constraints, feature representations, and internal noise. By explicitly linking behavioral performance to effective feature weighting and noise structure, this framework provides a foundation for future work aimed at establishing mechanistic links between neural architecture and perceptual performance across the visual field.

## Methods

### Ethics

All observers provided written informed consent and consented to the public release of their anonymized data. The experiment was conducted in accordance with the Declaration of Helsinki and was approved by the New York University ethics committee on activities involving human observers (IRB protocol number IRB-FY2022–6427).

### Participants

Twelve observers (8 females; age 22–30 years, mean ± SD: 26±2.82 years) participated. All but one observer (author SX) were naïve to the purpose of the study. All observers had normal or corrected-to-normal vision. Data collection was conducted between October 7, 2021 and March 28, 2024.

This study focused on polar-angle differences in contrast sensitivity, featural representation, and internal noise at 6° eccentricity. While a subset of these data was previously used to study eccentricity-dependent differences in visual performance [48], the present study addresses a different question: the computational basis of polar-angle asymmetries in vision. Ten of the twelve observers from that dataset were included here. Two observers were excluded from all analyses because their detection performance did not improve with decision variable at the upper vertical meridian, unlike the expected monotonic pattern shown in Figure 8A, left). This indicated that they were not reliably using task-relevant sensory information at that location. We then collected data from two additional observers. All analyses reported here are based on 12 observers.

### Apparatus

Stimuli were generated in MATLAB (MathWorks) using the Psychophysics Toolbox [83] and displayed on a gamma-linearized CRT monitor (Sony GDM-5402; 1280 × 960 pixels; 100 Hz; background luminance 33 cd*/*m^2^). Viewing distance was 57 cm, with head position stabilized by a chin rest. Eye position was recorded online with an EyeLink 1000 eye tracker positioned in front of the observer. Trials were aborted and repeated at the end of the block if gaze deviated by more than 1.5° of visual angle (dva) from fixation before stimulus offset.

### Experimental design

#### Stimuli and procedure

Figure 1B illustrates the trial sequence. Each trial began with a central black fixation cross (arm length = 0.3 dva) and five black placeholders (corner arm length = 0.75 dva), presented for 400 ms at the fovea and at four peripheral locations at 6° eccentricity: the left and right horizontal meridians and the lower and upper vertical meridians. Next, one target stimulus and four non-target stimuli (diameter = 3 dva) were presented for 100 ms. The target location was fixed within a block and varied across blocks. The target stimulus was either a signal Gabor embedded in filtered noise or filtered noise alone.

The signal Gabor, *G*(*x, y*), was a sinusoidal grating windowed by a Gaussian envelope:

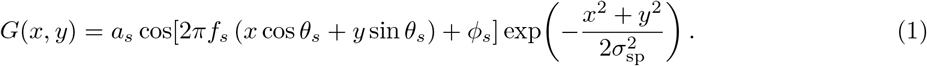

where *a*_*s*_ is the signal contrast, set by titration for each observer and location (see details below); *θ*_*s*_ is the signal orientation, which was always horizontal, *θ*_*s*_ = *π/*2; *f*_*s*_ is the signal spatial frequency, *f*_*s*_ = 2 cycles per degree (cpd); *ϕ*_*s*_ ∼ Unif(0, 2*π*) is a random phase drawn independently on each trial; and *σ*_sp_ is the standard deviation of the Gaussian envelope in the spatial domain. To enforce a fixed one-octave full-width at half-height (FWHH) in spatial frequency [84], we calculated *σ*_sp_ for each *f*_*s*_ (see derivation in Appendix A):

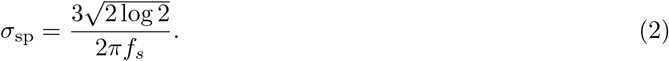

Filtered noise, *N*_*i*_, was generated independently on each trial *i* by bandpass-filtering white noise in the Fourier domain to retain spatial frequencies from 1 to 4 cpd with uniform power, followed by normalization to an RMS contrast of 20% (mean removed and standard deviation scaled appropriately). On signal-present trials, the target stimulus, 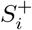, was the sum of the signal and noise, windowed by a circular aperture mask, *A*, with smoothed edges:

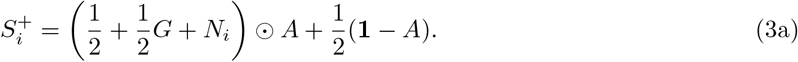

On signal-absent trials, the target stimulus was

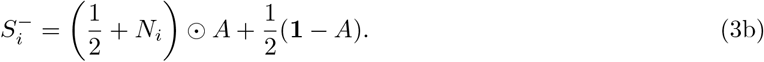

Here, ⊙ denotes element-wise multiplication, **1** is an all-ones matrix of matching size, and *S*_*i*_ is scaled from 0 to 1. This formulation makes the background outside the aperture equal to mid-gray luminance. Stimuli at non-target locations were noise patches generated independently on each trial and at each location following Equation 3b. In the following analyses, *S*_*i*_ denotes the target stimulus on trial *i*.

After a 500-ms post-stimulus delay, observers reported whether they perceived a Gabor in the target stimulus by pressing one of two keys (“F” for “present” and “J” for “absent”). Placeholders remained visible until response. Auditory feedback was provided after the response (high tone=correct; low tone=incorrect). Observers were instructed to prioritize accuracy, and there was no response deadline. To minimize spatial uncertainty, which increases with eccentricity [85] and may distort the recovered weight map in reverse-correlation analyses [86], the target location was kept constant within a block, and observers were informed of the target location at the start of each block.

### Titration and contrast threshold

To ensure that signal-absent trials provided informative, noise-driven response fluctuations, we anchored performance at a level that yielded enough false alarms for reverse correlation while minimizing response uncertainty or criterion instability. In this design, false alarms are critical because reporting a Gabor in pure noise indicates that trial-to-trial fluctuations in the noise can drive observers’ responses.

Before the main experiment, we used the Parameter Estimation by Sequential Testing (PEST) procedure [87] to target 70% accuracy, corresponding to a *d*^*′*^ ≈ 1 [55], separately for each observer and stimulus location. For each threshold estimate, we used two interleaved staircases of 50 trials each. The contrast at which performance converged to 70% accuracy served as the initial signal contrast for that location in the first block of the main experiment.

Because performance can improve with practice, we adjusted the signal contrast at each location beginning with its second block. This adjustment was based on the observer’s accuracy at that location in the preceding session: if accuracy was above or below the 70% target, contrast was decreased or increased proportionally, respectively. For example, 80% accuracy resulted in a 1% decrease in signal contrast.

The final contrast threshold for each location was defined as the geometric mean of the signal contrasts tested across sessions, computed by averaging log contrast and then transforming back to linear units. Contrast sensitivity was defined as the reciprocal of this threshold. This adaptive procedure maintained accuracy close to the 70% target across sessions, with an average deviation of approximately 6%. The mean deviation from the target across observers was 6.3% ± 0.3% at the left location, 6.4% ± 0.2% at the right location, 6.4% ± 0.3% at the lower location, and 6.6% ± 0.2% at the upper location.

Trials in the main experiment were organized into sessions, each containing five blocks—one per stimulus location—presented in randomized order. Across 10–15 one-hour sessions, observers completed an average of 4420 ± 160 trials per location, with equal numbers of trials at each location.

### Double-pass method and response consistency

To probe internal noise, we used a double-pass method coupled with the detection task to measure response consistency. Unknown to the observers, each block was divided into first and second halves, and each unique target stimulus presented in the first half was repeated in the second half. These matching trials were defined as pass 1 and pass 2 of the *p*-th pair. Distractor images were generated independently and therefore differed across passes.

Trial order was randomized within each half, ensuring that one member of each pair occurred in the first half of the block and the other in the second half. In a 100-trial block, 50 unique target stimuli were each presented twice, with equal numbers of signal-present and signal-absent trials in each pass. For each pair, observers could make either consistent responses, regardless of correctness, or inconsistent responses. Response consistency was calculated as the proportion of pairs yielding identical responses. Higher consistency reflects lower internal noise in the system [31]. We used response consistency as an empirical constraint when fitting the noisy-observer model.

### Psychophysical reverse correlation

To determine which stimulus features drove behavioral responses, we implemented psychophysical reverse correlation to estimate the perceptual weights assigned to a range of orientations and spatial frequencies [38–41, 43, 48]. This procedure comprised three steps (Figure 2A).

#### Step 1: Energy computation

For each trial *i*, we computed the contrast-energy profile, *E*_*i*_, of the target stimulus, *S*_*i*_, across orientation and spatial frequency:

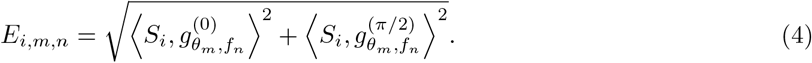

Here, ⟨·, ·⟩ denotes the inner product over pixels, and 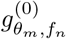 and 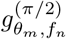 are quadrature-phase Gabor filters tuned to orientation *θ*_*m*_ and spatial frequency *f*_*n*_. Orientations spanned *θ*_*m*_ ∈ [−90°, 90°], with *m* = 1, …, *M* and *M* = 19, relative to the signal orientation. Spatial frequencies spanned *f*_*n*_ ∈ [1, 4] cpd, with *n* = 1, …, *N* (*N* = 19) and were sampled uniformly in log space. This energy computation approximates how V1 complex cells process visual information [88, 89]. Higher energy indicates stronger feature structure in the corresponding orientation–spatial-frequency channel. Importantly, even when no signal was present, filtered noise could yield high energy near the signal orientation and spatial frequency, producing false-alarm responses.

For each (*θ*_*m*_, *f*_*n*_) pair, we computed the mean and standard deviation across all signal-absent trials and defined 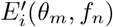 as the *z*-scored value of *E*_*i*_(*θ*_*m*_, *f*_*n*_). The same normalization was applied to signal-present and signal-absent trials.

#### Step 2: Weight estimation

After computing the trial-wise energy profiles, we estimated how fluctuations in orientation–spatial-frequency energy predicted observers’ binary responses. We fitted a multivariate binomial generalized linear model with a probit link to signal-absent trials, using the normalized energy profile from each trial to predict the probability of a “present” response:

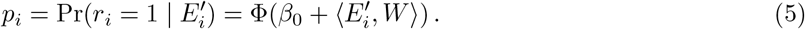

where *r*_*i*_ ∈ {0, 1} is the observed binary response, with *r*_*i*_ = 1 denoting a “present” response and *r*_*i*_ = 0 denoting an “absent” response. Φ(·) is the standard normal cumulative distribution function, *β*_0_ is the intercept, which was excluded from subsequent analyses, and *W* is the estimated two-dimensional weight map.

We estimated *W* using only signal-absent trials because, in the absence of a physical signal, noise-driven fluctuations are most diagnostic of the weights assigned to task-relevant features [35, 90–92].

To stabilize the reverse-correlation estimates, we represented *W* using smooth separable basis functions (Appendix B). Orientation weights were modeled as a weighted sum of three von Mises basis functions, and spatial-frequency weights were modeled as a weighted sum of three asymmetric Gaussian basis functions. This basis configuration was selected using model simulations that evaluated recovery of the orientation and spatial-frequency tuning functions.

#### Step 3: One-dimensional tuning-function characterization

After estimating the two-dimensional weight map, we marginalized it along each feature dimension to obtain one-dimensional weights, denoted *w*_*θ*_ and *w*_*f*_, respectively. These marginalized weights were then fitted with descriptive tuning functions to extract summary measures of sensitivity and selectivity. These post hoc fits were used only to characterize the recovered weight map; they were not used to estimate the two-dimensional template or to fit the noisy-observer model.

The marginalized orientation weights were fitted with a von Mises function:

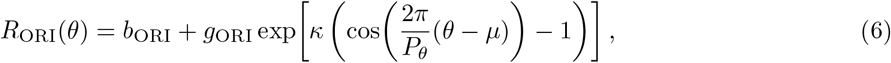

where *b*_ORI_ is the baseline, *g*_ORI_ is the gain above baseline, *µ* is the preferred orientation (fixed at 0°), *κ* controls concentration, and *P*_*θ*_ = 180° is the period of orientation space. To express orientation bandwidth in interpretable units (degrees), we quantified it as the full-width at half of the fitted peak-to-trough height. We first calculated the half-height of the fitted function:

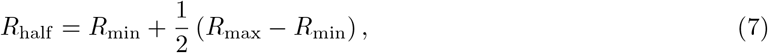

where *R*_max_ and *R*_min_ are the maximum (peak) and minimum (trough) of the fitted curve. We then numerically identified the first crossing to the right of the peak, *θ*_half_, at which *R*_ORI_(*θ*_half_ ) = *R*_half_ . Because the von Mises function is symmetric about its peak, orientation bandwidth (full width at half-height) was calculated as twice the half-width at half-height.

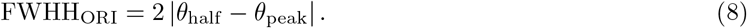

The marginalized spatial-frequency weights were fitted with a log-Gaussian function:

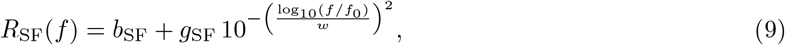

where *b*_SF_ is the baseline, *g*_SF_ is the gain above baseline, *f*_0_ is the peak spatial frequency, and *w* controls bandwidth in log-frequency space. Spatial-frequency bandwidth was calculated at the half-height defined in Equation 7. We identified the left and right spatial frequencies, *f*_*L*_ and *f*_*R*_, at which the fitted curve crossed this half-height. The full spatial-frequency bandwidth was then expressed in octaves:

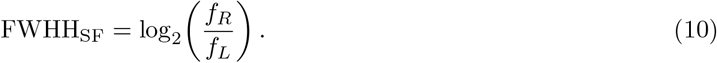

### Noisy-observer model

To jointly account for trial-wise detection responses and double-pass consistency, we constructed a noisy-observer model (Figure 2B). For each observer and stimulus location, the model used the perceptual template—the two-dimensional weight map *W* estimated via reverse correlation—to predict trial-wise behavioral responses.

The initial decision variable was

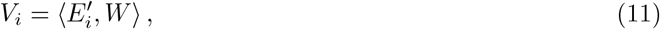

which indexes the amount of task-relevant information in the stimulus on trial *i*. Larger values of *V*_*i*_ imply a higher probability of reporting that the signal was present.

To place the decision variable on a normalized scale, such that a given amount of decision-stage noise had a comparable influence across observers and locations, we divided *V*_*i*_ by its standard deviation across all signal-present and signal-absent trials, *i* = 1, …, *T*, yielding

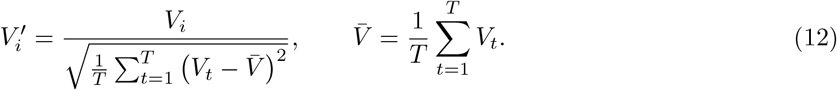

The observer’s final decision on trial *i* depended on a noisy decision variable, 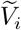, normally distributed around 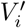 with variance 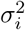:

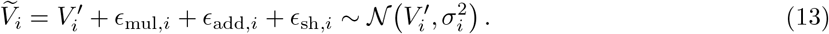

Total internal noise comprised three Gaussian components. Multiplicative noise was private to each pass and scaled with stimulus strength: 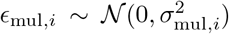, where 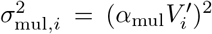. Additive noise was private to each pass and constant across trials, with 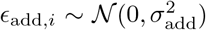. Shared noise was common to the two passes of a double-pass pair, with 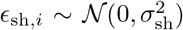. This shared component represents a trial-locked fluctuation associated with the specific physical stimulus. It is not correlated with the template response but is perfectly correlated across repeated presentations of the same noise field. Thus, if double-pass pair *p* contains trials *i* and *j*, then *ϵ*_sh,*i*_ = *ϵ*_sh,*j*_.

The total marginal noise variance on trial *i* was therefore

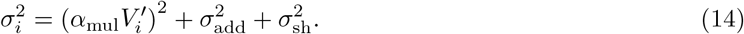

The model generated two behavioral probabilities. On a given trial, the observer responded “present” if 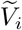 exceeded a decision criterion *k*. The model therefore predicted the marginal detection probability

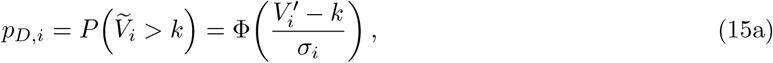

where Φ(·) is the standard normal cumulative distribution function.

For each double-pass pair *p*, the consistency probability was the probability of producing the same response on both passes:

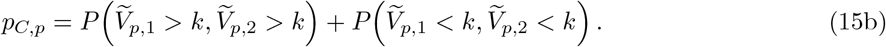

To capture shared fluctuations between passes, as commonly assumed in models of double-pass data [52, 93, 94], we modeled 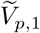 and 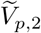 jointly as a bivariate normal distribution:

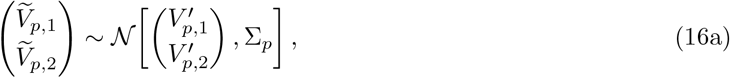

with covariance matrix

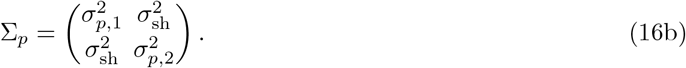

Setting *σ*_sh_ = 0 is equivalent to assuming independent internal noise across the two passes. Under independence, the expected consistency probability for a marginal detection probability *p*_*D,i*_ is 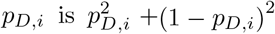, which was used as the baseline in the binned plots.

Jointly modeling detection and consistency probabilities helps dissociate private and shared noise. Detection probability depends on the total marginal variance, whereas the relative amount of shared and private noise affects the joint consistency probability. Thus, detection responses constrain the trial-wise marginal noise level and decision criterion, whereas consistency responses constrain shared covariance relative to private noise.

We fitted the free parameters *ψ* = {*α*_mul_, *σ*_add_, *σ*_sh_, *k*} by maximizing the joint log-likelihood of two binary datasets. For trial-wise detection responses, *d*_*i*_ ∈ {0, 1}, with *d*_*i*_ = 1 denoting a “present” response,

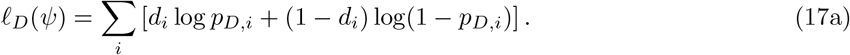

For pair-wise consistency responses, *c*_*p*_ ∈ {0, 1}, with *c*_*p*_ = 1 denoting a consistent response pair,

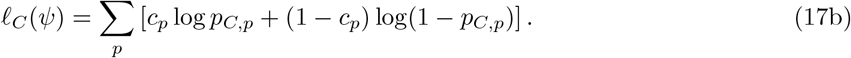

where *p*_*D,i*_ and *p*_*C,p*_ are the model-predicted probabilities of detection on trial *i* and consistency for pair *p*, respectively. We optimized *ℓ*_*D*_(*ψ*) + *ℓ*_*C*_(*ψ*) using Bayesian adaptive direct search (BADS [95]). BADS explores the parameter space by alternating between fast local Bayesian-optimization steps and a systematic, nonlocal exploration of a mesh grid.

### Monte Carlo cross-validation

We evaluated the model using Monte Carlo cross-validation. On each iteration, we randomly partitioned trials into three non-overlapping sets—template, training, and held-out test—while keeping double-pass pairs intact within each set. We derived the template from the template set using 80% of the signal-absent trials, equivalent to 40% of all trials. From the remaining trials, we constructed a training set comprising 15% of all trials by sampling 15% of the remaining signal-absent trials and 15% of the signal-present trials. We then constructed a test set comprising 5% of all trials, including 5% of the remaining signal-absent trials and 5% of the signal-present trials, with no overlap with the training set. We repeated this cross-validation procedure for 1000 iterations per location and observer and reported the medians of the parameter estimates and model predictions.

To summarize predicted performance as a function of stimulus strength, we divided held-out test trials into six quantiles of the standardized decision variable, 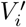. Quantile boundaries were computed once per observer and location using all trials before cross-validation. Within each bin, we calculated the empirical detection rate, defined as the proportion of “present” responses, and the empirical response consistency, defined as the proportion of identical responses. We also averaged the model-predicted detection and consistency probabilities across held-out test trials within each bin.

Likelihood calculations did not depend on binning, and the primary modeling results were insensitive to the number of bins. For descriptive goodness of fit, we calculated the root-mean-square error (RMSE) within each observer and location and then averaged these values across observers.

To evaluate the structural role of the data-derived feature representation and the contribution of each noise parameter, we fitted nested model variants: a full model containing all three noise parameters, three models that each excluded one noise parameter, and three models that each included only one noise parameter. We compared these variants using ΔNLL = NLL_model_ − min(NLL), where the minimum was calculated across all variants fitted at a given stimulus location. Each model variant was fitted independently within each cross-validation split.

### Model simulation

To validate the recovery pipeline, we conducted simulations in which responses were generated from known feature templates, internal-noise parameters, and criterion values. We then applied the full analysis pipeline—including template estimation and noisy-observer model fitting—to the simulated data.

Recovery was evaluated for the two-dimensional template, marginalized orientation and spatial-frequency tuning functions, model parameters, and model identity. Full simulation details and recovery results are reported in Appendix C. These simulations were used to select the basis-function hyperparameter for the empirical analyses and to identify model variants with weak parameter interpretability.

The simulations showed that the template-estimation procedure reliably recovered the ground-truth feature template, particularly the marginalized orientation tuning function (Figure S2). Parameter- and model-recovery analyses further indicated that most reduced models were recoverable (Figure S5), but also revealed one limitation: the full model, which included both multiplicative and additive noise, showed a tradeoff between these parameters (Figure S6), making their individual estimates difficult to interpret.

### Statistical analysis

#### Combining locations

To quantify polar-angle asymmetries—the horizontal–vertical anisotropy (HVA) and vertical-meridian asymmetry (VMA)—we performed planned comparisons on combined metrics within each observer. These metrics included behavioral measures (contrast sensitivity, response consistency, *d*^*′*^, accuracy, criterion, and response time), reverse-correlation template measures (two-dimensional weights, marginalized tuning functions, and tuning parameters), noisy-observer model predictions (predicted detection and consistency probabilities), and model estimates (internal-noise parameters).

The horizontal meridian was defined as the mean of the metric at the left and right locations, and the vertical meridian was defined as the mean at the upper and lower locations. This approach ensured that all single and combined locations contained the same number of trials.

Asymmetry magnitude was quantified using the normalized location-difference index 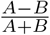, where *A* and *B* are the metrics at the two locations being compared. For the HVA, *A* was the horizontal meridian and *B* was the vertical meridian. For the VMA, *A* was the lower vertical meridian and *B* was the upper vertical meridian.

#### Cross-validation and single-observer summaries

Behavioral performance was computed from all trials, yielding one value per observer and location without resampling. Reverse-correlation and model-derived outputs were estimated using Monte Carlo cross-validation. For each observer and location, cross-validated measures were summarized by their median across iterations. When plotting single-observer estimates, error bars denote the 68% confidence interval across cross-validation iterations. These observer-level medians were carried forward to the group-level analyses.

#### Bootstrap uncertainty and effect sizes

Group-level summaries and uncertainty were obtained by bootstrap resampling observers 10,000 times. On each resample, observers were drawn with replacement and the statistic of interest was recomputed, yielding a bootstrap distribution. We report the bootstrapped median and confidence intervals, using 95% confidence intervals for inference and 68% confidence intervals for plotted group error bars.

Throughout the Results, we quantified effects using five complementary analyses:

1. Factor effects were summarized using one-or two-way repeated-measures ANOVAs, reporting partial eta-squared, 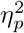, as the effect size with bootstrapped confidence intervals. When location was included as a factor, it had three levels: horizontal meridian, lower vertical meridian, and upper vertical meridian. Permutation *p*-values were obtained from 10,000 permutations. On each permutation, condition labels were shuffled within observers to generate a null distribution of *F* statistics. The permutation *p*-value was the proportion of permuted *F* values greater than or equal to the observed *F* value.
2. Planned location contrasts were summarized using paired differences for HM–VM and LVM–UVM comparisons. We report the bootstrapped median and 95% confidence interval of the group-mean paired difference, together with Hedges’ *g* as the standardized effect size. Permutation *p*-values were obtained from 10,000 random sign flips of the within-observer difference scores. The permutation *p*-value was the proportion of permuted |*t*| values greater than or equal to the observed |*t*|.
3. Correlations between contrast sensitivity and template-related or model-derived estimates across observers were summarized using partial Pearson correlations controlling for visual-field location. We report Pearson’s *r* with bootstrapped confidence intervals. Two-tailed permutation *p*-values were calculated by shuffling observer labels and computing the proportion of permuted |*r*| values greater than or equal to the observed |*r*|.
4. Correlations between asymmetries (HVA or VMA) across observers were assessed by correlating the location-difference indices separately for the HVA and VMA. We reported Spearman’s *ρ*, which is less sensitive to influential observations and does not require a linear relationship, an important consideration given the small sample size. When the expected direction was specified a priori based on analysis 3, we report two-tailed 90% bootstrap confidence intervals and one-tailed *p*-values. Otherwise, we report two-tailed 95% bootstrap confidence intervals and two-tailed *p*-values. Permutation *p*-values were calculated as described for analysis 3.
5. To ensure computational reproducibility, all bootstrap and permutation procedures were run using fixed random seeds.

## Appendix A Deriving the Gaussian-envelope SD from the desired spatial-frequency bandwidth

To set the spatial-frequency bandwidth of the Gabor filters to a desired value—one octave in the present study—we derived the spatial-domain standard deviation, *σ*_sp_, used in Equation 2. The Fourier transform of a Gabor with preferred spatial frequency *f*_*n*_ contains two Gaussian components centered at ±*f*_*n*_. Considering the positive-frequency component, its response as a function of spatial frequency *f* is

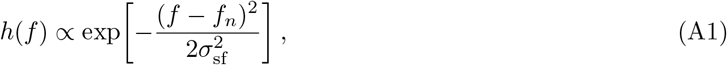

where *σ*_sf_ is the standard deviation in the spatial-frequency domain.

The frequencies at which the response falls to one-half of its peak satisfy

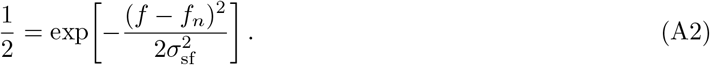

Solving for *f* gives

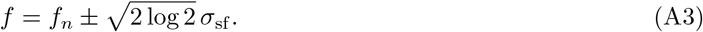

For a one-octave bandwidth, the ratio of the upper and lower half-height frequencies must equal 2:

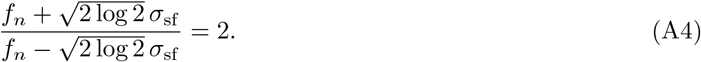

Solving for *σ*_sf_ yields

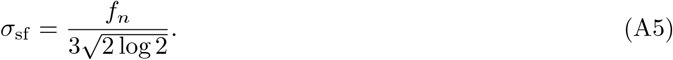

For a Gaussian, the spatial-domain and spatial-frequency-domain standard deviations are inversely related:

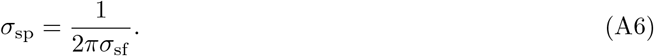

Substituting Equation A5 into Equation A6 gives

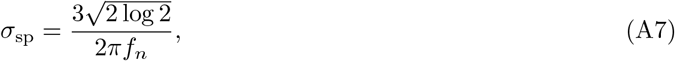

which corresponds to Equation 2. Thus, to maintain a one-octave spatial-frequency bandwidth, the spatial standard deviation of the Gaussian envelope scales inversely with the filter’s preferred spatial frequency.

## Appendix B Basis-function parameterization and hyperparameter selection

From Equation 4, we computed the energy matrix on trial *i, E*_*i*_ ∈ ℝ^*M ×N*^, sampled over *M* orientation channels and *N* spatial-frequency channels. To impose smoothness on the estimated perceptual template, we projected each normalized energy matrix, 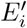, onto low-dimensional sets of orientation and spatial-frequency basis functions.

Let 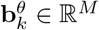, *k* = 1, …, *K*, denote the *k*th orientation basis vector evaluated across the *M* orientation channels, and let 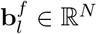, *l* = 1, …, *L*, denote the *l*th spatial-frequency basis vector evaluated across the *N* spatial-frequency channels. We collected these vectors as columns of the basis matrices 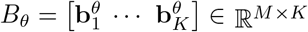 and 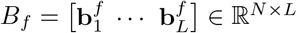.

We projected the trial-wise normalized energy matrix onto the separable basis:

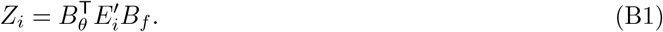

The resulting matrix *Z*_*i*_ ∈ ℝ^*K×L*^ contains the basis-space coefficients for trial *i* and serves as the predictor matrix for the generalized linear model. Equivalently, the coefficient associated with the *k*th orientation basis function and the *l*th spatial-frequency basis function is 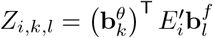.

Because neighboring basis functions respond to similar orientations and spatial frequencies, the projected coefficients can be correlated across trials. We therefore used ridge regularization to stabilize coefficient estimation.

Before fitting the generalized linear model, each projected coefficient was standardized across trials:

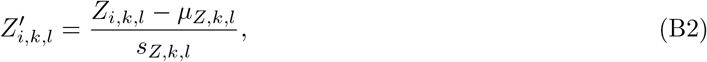

where *µ*_*Z,k,l*_ and *s*_*Z,k,l*_ are the across-trial mean and standard deviation of *Z*_*i,k,l*_, respectively. This standardization placed all projected coefficients on a common scale, allowing the ridge penalty to act comparably across basis components.

Using the standardized projected coefficients as predictors, the probit generalized linear model corresponding to Equation 5 was

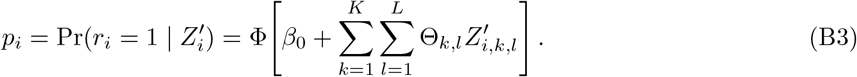

Here, Φ(·) is the standard normal cumulative distribution function, *β*_0_ is the intercept, and Θ_*k,l*_ is the regression coefficient associated with the standardized predictor 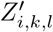. Positive values of Θ_*k,l*_ indicate that greater energy aligned with the corresponding orientation–spatial-frequency basis pattern increases the probability of a “present” response, whereas negative values indicate the opposite.

We estimated the coefficients by maximizing the ridge-regularized log-likelihood

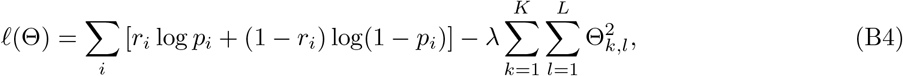

where *λ* controls the strength of regularization. Larger values of *λ* shrink the fitted coefficients toward zero and reduce estimation variance. Smoothness is imposed primarily by the low-dimensional basis representation, whereas ridge regularization stabilizes estimation within that basis.

Because the generalized linear model was fitted using standardized projected coefficients, Θ_*k,l*_ describes response modulation per one-standard-deviation increase in *Z*_*i,k,l*_. Before reconstructing the template in the original orientation-by-spatial-frequency channel space, we converted the coefficients back to the unstandardized projected-energy scale:

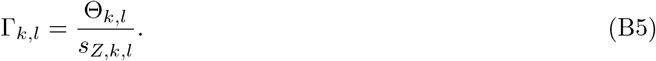

Collecting these coefficients across all basis-function pairs gives Γ ∈ ℝ^*K×L*^. The smooth two-dimensional weight map was reconstructed as

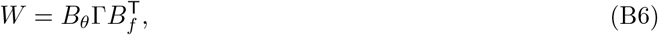

where *W* ∈ ℝ^*M ×N*^ matches the dimensions of the original energy matrix.

### Orientation basis functions

We evaluated each orientation basis function at every sampled orientation channel *θ*_*m*_, where *m* = 1, …, *M* . The orientation basis centers, 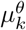, were evenly spaced across one orientation period, *P*_*θ*_ = 180°, and included one basis centered on the target orientation:

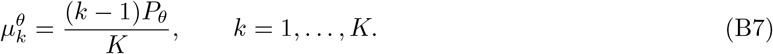

The circular distance between orientation *θ*_*m*_ and the *k*th basis center was

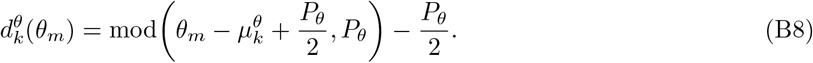

where mod(*a, b*) returns the remainder after division of *a* by *b*. Equation B8 wraps orientation differences onto the interval 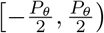, so that the basis center is mapped to zero and all other orientations lie within ±90° of it.

The value of the *k*th von Mises basis function at channel *θ*_*m*_ was

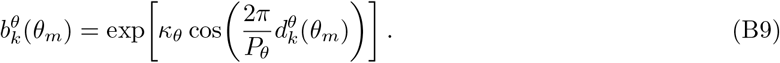

Evaluating this function across all *M* sampled orientation channels gives the unnormalized basis vector

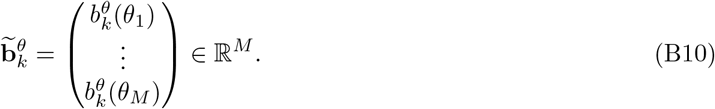

The concentration parameter *κ*_*θ*_ was defined from the bandwidth parameter *σ*_*θ*_:

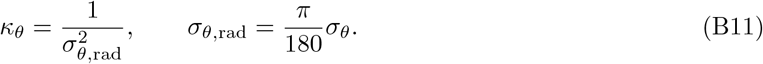

Both *K* and *σ*_*θ*_ were treated as hyperparameters.

### Spatial-frequency basis functions

The spatial-frequency basis centers, 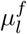, were evenly spaced across the sampled log-spatial-frequency range:

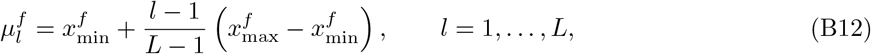

Where 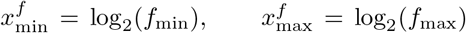. The signed distance between the *n*th spatial-frequency channel and the *l*th basis center was

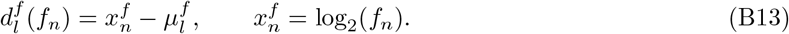

We used an asymmetric Gaussian basis family because pilot analyses indicated that asymmetric Gaussians described the spatial-frequency profiles better than symmetric log-Gaussians. The value of the *l*th spatial-frequency basis function at channel *f*_*n*_ was

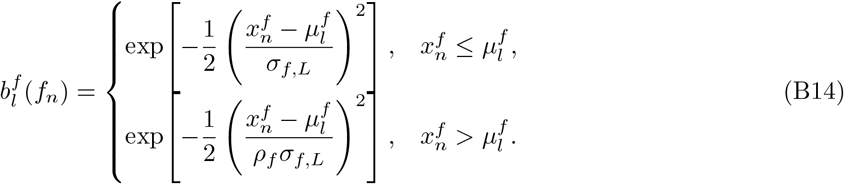

Here, *σ*_*f,L*_ is the left-side bandwidth and *ρ*_*f*_ is the ratio between the right- and left-side bandwidths. Evaluating this function across all *N* sampled spatial-frequency channels gives the unnormalized basis vector

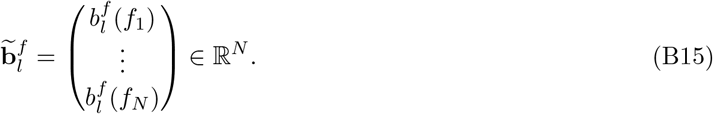

The hyperparameters defining the spatial-frequency basis set were *L, σ*_*f,L*_, and *ρ*_*f*_ .

### Basis-vector normalization

Before projection, each orientation and spatial-frequency basis vector was normalized to unit Euclidean norm so that basis functions with different bandwidths had comparable total magnitude:

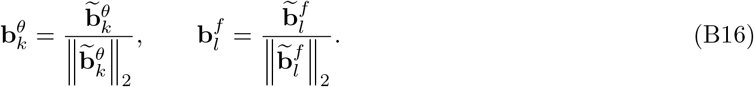

The normalized vectors 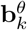 and 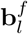 were used to construct the basis matrices in Equation B1.

### Hyperparameter selection

Hyperparameters for template estimation were selected using five-fold cross-validation. For each candidate hyperparameter combination, we fitted the ridge-regularized binomial generalized linear model to four folds and evaluated the negative log-likelihood on the held-out fold. The candidate grid included *K* ∈ {3, …, 8}, *L* ∈ {3, …, 8}, *σ*_*θ*_ ∈ {0.5, 0.6, …, 0.9}, *σ*_*f,L*_ ∈ {0.4, 0.5, …, 0.9}, *ρ*_*f*_ ∈ {0.7, 0.9, 1.1, 1.3, 1.5}, and *λ* ∈ {10^−3^, 10^−2^, 10^−1^, 1, 10, 10^2^, 10^3^}. To summarize the selection procedure, we calculated the frequency with which each candidate value was selected across iterations and simulation conditions. The simulation conditions varied signal contrast *a*_*s*_, SDT criterion *k*_SDT_, multiplicative noise *α*_mul_, additive noise *σ*_add_, and shared noise *σ*_sh_. The most consistently selected configuration was *K* = 3, *L* = 3, *σ*_*θ*_ = 0.6, *σ*_*f,L*_ = 0.4, *ρ*_*f*_ = 0.9, and *λ* = 100. We therefore used this configuration for all subsequent template-estimation and noisy-observer-model analyses.

## Appendix C Model simulation and recovery validation

To validate the recovery pipeline, we simulated datasets with known ground-truth templates, internal-noise parameters, decision criteria, and model structures. We then applied the complete analysis pipeline used for the empirical data, including reverse-correlation template estimation and noisy-observer-model fitting, and evaluated recovery of the template, model parameters, and generating model.

On each simulated trial, bandpass-filtered external noise was sampled with RMS contrast fixed at 20%, matching the experimental stimuli. Signal presence was assigned independently across trials. Each stimulus was transformed into an orientation-by-spatial-frequency energy profile using the same energy-computation procedure as in Equation 4. The noiseless decision variable was computed as the inner product of this energy profile and the ground-truth template, as in Equation 13, and standardized as in Equation 14.

Internal noise was then added to generate the noisy decision variable used to produce simulated responses, following Equation **??**. When present in the generating model, multiplicative and additive noise were sampled independently on each trial, whereas shared noise was sampled once per double-pass pair and applied identically to both passes. A binary response was generated by comparing the noisy decision variable with a decision criterion.

### Converting the criterion from SDT units to decision-variable units

In the simulations, we specified the decision criterion in standard signal-detection-theory units because this scale is interpretable across simulation conditions. However, the noisy-observer model generates responses by comparing the noisy decision variable with a criterion expressed in decision-variable units. For each simulation condition, we therefore converted the desired SDT criterion, *k*_SDT_, to a decision-variable criterion, *k*.

For a candidate value of *k*, we calculated the model-implied hit rate, *P* (*r* = 1 | *S*^+^; *k*), and false-alarm rate, *P* (*r* = 1 | *S*^−^; *k*), where *S*^+^ denotes signal-present trials, *S*^−^ denotes signal-absent trials, and *r* = 1 denotes a simulated “present” response. These probabilities determine the SDT criterion implied by *k*:

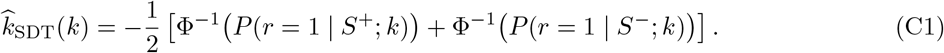

We then selected the decision-variable criterion whose implied SDT criterion was closest to the desired value:

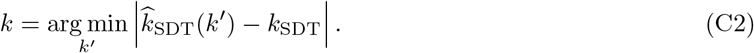

This procedure ensured that simulated observers had the intended response bias in SDT units, even though responses were generated in decision-variable units.

After generating binary responses, we applied the same analysis pipeline used for the empirical data. The template was re-estimated from simulated responses using reverse correlation, and the noisy-observer model was fitted to recover the internal-noise parameters and decision criterion. For each simulation condition, we simulated 4000 trials and repeated the recovery procedure for 1000 iterations.

### Candidate model configurations

We simulated datasets under seven generating models. The full model included all three noise parameters: *α*_mul_, *σ*_add_, *σ*_sh_. The six reduced models were obtained by fixing one or more noise parameters to zero:

1. M1, FullModel: multiplicative, additive, and shared noise;
2. M2, NoMul: additive and shared noise;
3. M3, NoAdd: multiplicative and shared noise;
4. M4, NoShared: multiplicative and additive noise;
5. M5, MulOnly: multiplicative noise only;
6. M6, AddOnly: additive noise only;
7. M7, SharedOnly: shared noise only.

In every model, the decision criterion *k* was included as a free parameter. Data generated by each of the seven models were fitted with all seven candidate models, yielding 49 generating-model–fitted-model combinations.

### Ground-truth parameter sampling and simulation-condition composition

Ground-truth internal-noise parameters were sampled from ranges chosen to produce behaviorally plausible performance: *α*_mul_ ∈ {0.2, 0.4, 0.6, 0.8, 1.0}, *σ*_add_ ∈ {0.5, 1, 1.5, 2, 2.5}, and *σ*_sh_ ∈ {0.5, 1, 1.5, 2, 2.5}. Signal contrast varied over *a*_*s*_ ∈ {0.3, 0.5}, covering the range observed in the empirical data. The true criterion was specified in SDT units: *k*_SDT_ ∈ {−0.2, 0, 0.2}, and was then converted to decision-variable units using Equation C2. This range also covered the criteria observed in the empirical data.

Each unique combination of generating model, signal contrast, SDT criterion, and applicable internal-noise values constituted one simulation condition, yielding 1,290 conditions. To focus on behaviorally relevant regimes, we retained only conditions in which simulated percent correct was between 60% and 80%. This filter retained 655 conditions, whose composition is summarized in Figure S1.

**Figure S1.**
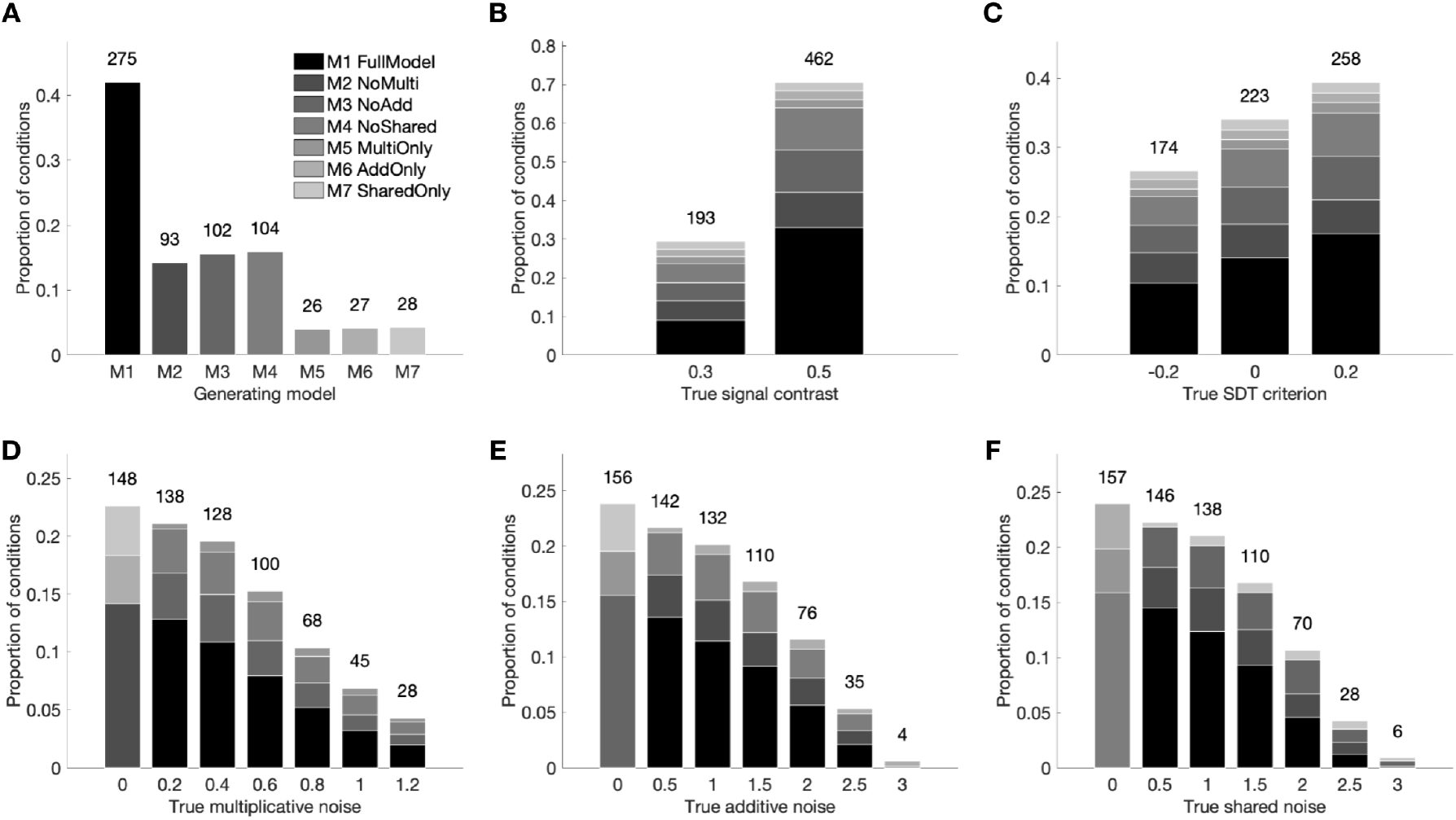
Composition of simulation conditions retained after the accuracy filter. Each panel shows the distribution of retained simulation conditions as a function of one ground-truth factor: **(A)** generating model, **(B)** signal contrast, **(C)** SDT criterion, **(D)** multiplicative noise, **(E)** additive noise, and **(F)** shared noise. Conditions were retained when simulated percent correct was between 60% and 80%. Bar shading denotes generating-model identity, as indicated by the grayscale legend in Panel A. Retained conditions were broadly distributed across signal contrast, SDT criterion, and internal-noise values, indicating that the filter did not strongly concentrate the retained set at a single parameter value. The largest imbalance occurred across generating models: the full model contributed the most retained conditions, whereas reduced models contributed fewer because one or more noise parameters were fixed to zero. The full model spans a broader range of decision-variable distributions and therefore contains more parameter combinations capable of producing intermediate performance. Thus, its overrepresentation follows from the performance-based inclusion criterion rather than preferential sampling.

### Template recovery

Figure S2 summarizes recovery of the ground-truth template across internal-noise parameters, signal contrast, and criterion. Recovery worsened as internal noise increased, indicating that noisier decision processes reduced the information available for estimating the template. Recovery also varied with signal contrast but showed little systematic dependence on the SDT criterion.

**Figure S2.**
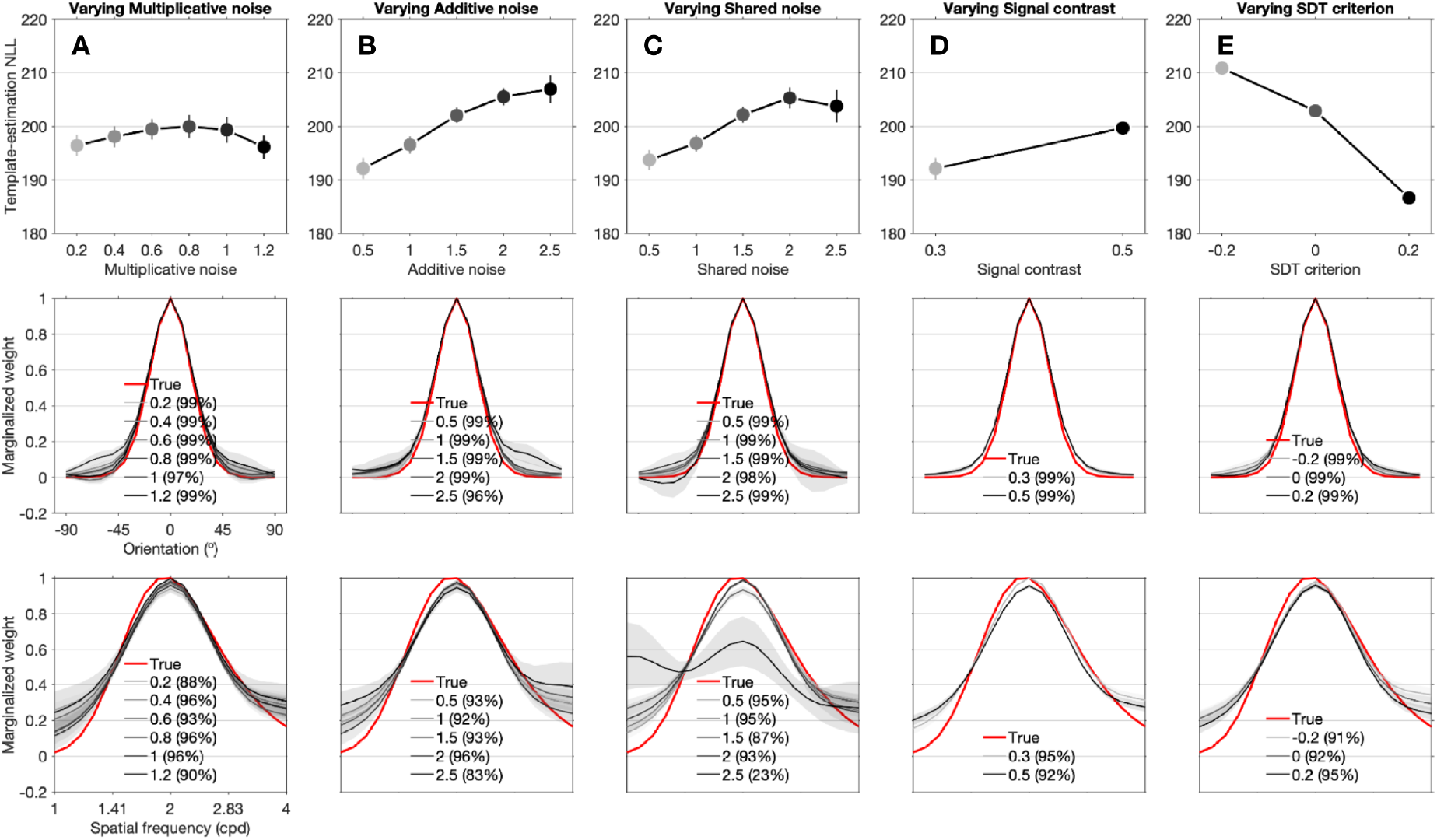
Template recovery across ground-truth parameters. Each column corresponds to one varied ground-truth factor, from left to right: multiplicative noise, additive noise, shared noise, signal contrast, and SDT criterion. Top row: negative log-likelihood (NLL) of template estimation as a function of the varied ground-truth factor, averaged across the relevant simulation conditions. Error bars denote *±*1 SEM; lower NLL indicates better model fit. Dot shading indicates the value of the varied factor, and the same shading scheme is used throughout the figure. Middle row: marginalized orientation tuning functions for the ground-truth template (red) and recovered templates at different values of the varied factor. Shaded bands denote *±*1 SEM. The legend in each panel reports the ground-truth factor value and the coefficient of determination, *R*^2^, between the ground-truth and recovered marginalized tuning functions. Bottom row: marginalized spatial-frequency tuning functions, plotted using the same conventions as in the middle row.

Even when recovery of the complete two-dimensional template was degraded, the marginalized tuning functions remained close to the ground truth, particularly in the orientation dimension. Spatial-frequency recovery was less precise. These simulations therefore support the ability of reverse correlation to recover task-relevant tuning structure while motivating caution when interpreting fine-grained spatial-frequency estimates.

### Behavioral-metric recovery

Figure S3 shows recovery of detection and response consistencies across generating and fitted models. Open circles show metrics calculated from data simulated using the ground-truth parameters and model structures after trials were divided into six decision-variable bins. Binning was required because these metrics are defined over groups of trials.

In the top row, open circles show detection rate, defined as the proportion of “present” responses within each bin. The corresponding curves show the trial-wise detection probabilities from Equation 17a, averaged within each bin. In the bottom row, open circles show response consistency, defined as the proportion of double-pass pairs with identical responses, and curves show the corresponding predicted consistency probabilities from Equation 17b.

For most generating models, the matched fitted model captured both metrics well. Response consistency was especially diagnostic because model predictions differed more strongly across fitted models than did predictions of detection rate.

### Parameter recovery

Figure S4 shows parameter recovery for matched generating–fitted model pairs. From top to bottom, the rows compare recovered and ground-truth values of multiplicative noise, additive noise, shared noise, and criterion in decision-variable units.

Overall, recovered estimates followed the identity line, indicating successful parameter recovery across several generating models. Recovery was generally robust to variation in internal noise, signal contrast, and criterion, although precision differed among parameters and model variants. A notable exception was the tradeoff between multiplicative and additive noise in the FullModel and NoShared models. In both models, multiplicative noise tended to be underestimated and additive noise overestimated, motivating the parameter-correlation analysis reported in Figure S6.

### Model recovery

To assess whether the fitting procedure identified the model structure that generated the data, we examined how often each candidate model provided the best fit to data generated by each model (Figure S5). The generating model was often favored, as indicated by elevated win rates along the diagonal of the generating-model-by-fitted-model matrix.

Recovery was not perfectly selective. Closely related nested models often produced similar negative log-likelihoods and comparable predictions of detection rate, accuracy, and response consistency. Thus, multiple model structures could reproduce the same behavioral summaries with similar precision.

Nevertheless, the simulations revealed informative structure. Models containing shared noise were distinguishable from models that omitted it because shared noise altered the dependence between responses across repeated trials. Parameter recovery was accurate for several reduced models but deteriorated when fitted models were overparameterized or misspecified. Thus, good behavioral fit did not necessarily guarantee precise recovery of individual noise components.

**Figure S3.**
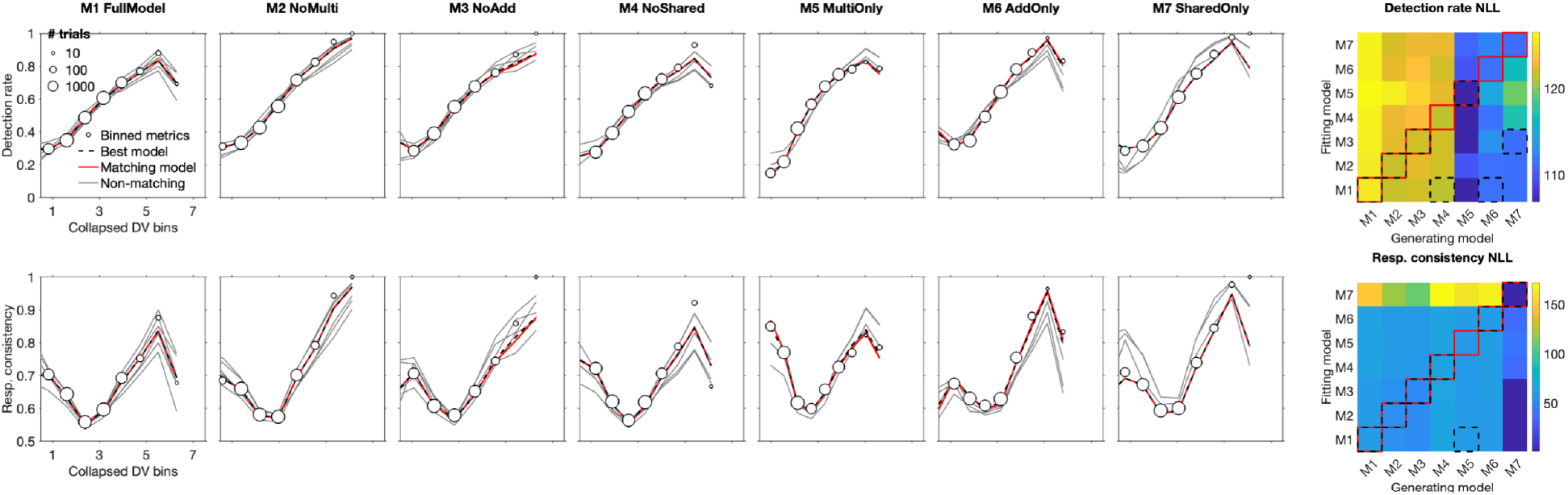
Recovery of binned behavioral metrics across generating and fitted models. Rows correspond to detection rate and response consistency. The rightmost column shows heatmaps of RMSE between simulated and model-predicted metrics for each fitted model (rows) and generating model (columns), averaged across included simulation conditions. Darker cells indicate lower RMSE. Solid red boxes indicate matched generating–fitted model pairs on the diagonal, and dashed black boxes indicate the fitted model with the lowest RMSE within each generating-model column. The remaining columns show binned behavioral summaries for each generating model (M1–M7), averaged across included simulation conditions. In each panel, the horizontal axis shows decision-variable bins. Black circles show metrics calculated from simulated binary responses, with circle size proportional to the number of trials in each bin. Curves show predicted detection or consistency probabilities averaged within each bin. The red curve indicates the matched fitted model, gray curves indicate mismatched fitted models, and the dashed black curve indicates the fitted model with the lowest RMSE for that generating model and metric. When the red and black indicators overlap, the matched model also provided the best recovery. RMSE values were weighted by the number of trials contributing to each bin.

Overall, the simulations indicate that the modeling framework can recover broad features of the underlying noise structure, whereas exact model identity and individual parameter estimates should be interpreted cautiously for closely related nested models.

### Parameter correlations

Figure S6 shows pairwise correlations between recovered parameters across fitting iterations for matched generating–fitted models. These correlations reveal dependencies within the fitted parameter space.

Because correlations in models containing several noise parameters may reflect interactions among multiple components, we focused on correlations between pairs of noise parameters. Multiplicative and additive noise showed a negative correlation, indicating a tradeoff during estimation. The other noise-parameter pairs showed positive correlations. The tradeoff between multiplicative and additive noise indicates that models containing both components are difficult to interpret because changes in one parameter can be partially offset by changes in the other.

**Figure S4.**
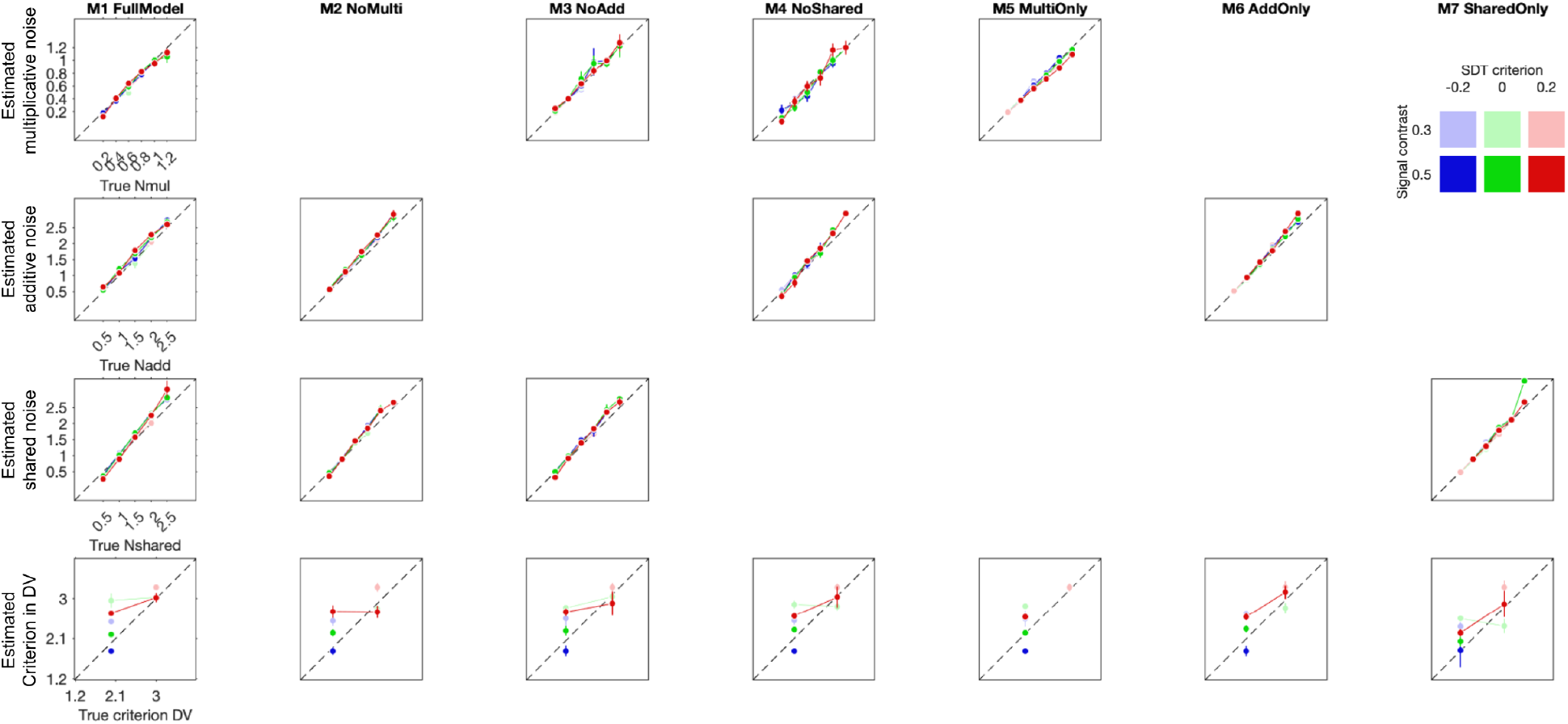
Parameter recovery. Each panel plots recovered parameter values against ground-truth values. Rows correspond to multiplicative noise, additive noise, shared noise, and criterion in decision-variable units. Columns correspond to matched generating–fitted model pairs. Each point summarizes one signal-contrast *×* SDT-criterion condition after averaging across the remaining simulation conditions; vertical error bars denote *±*1 SEM. Color encodes condition identity, with hue indicating SDT criterion and lightness indicating signal contrast. The dashed identity line indicates perfect recovery. For visualization, true decision-criterion values were binned before plotting. Panels are omitted when a parameter was absent from the corresponding model.

**Figure S5.**
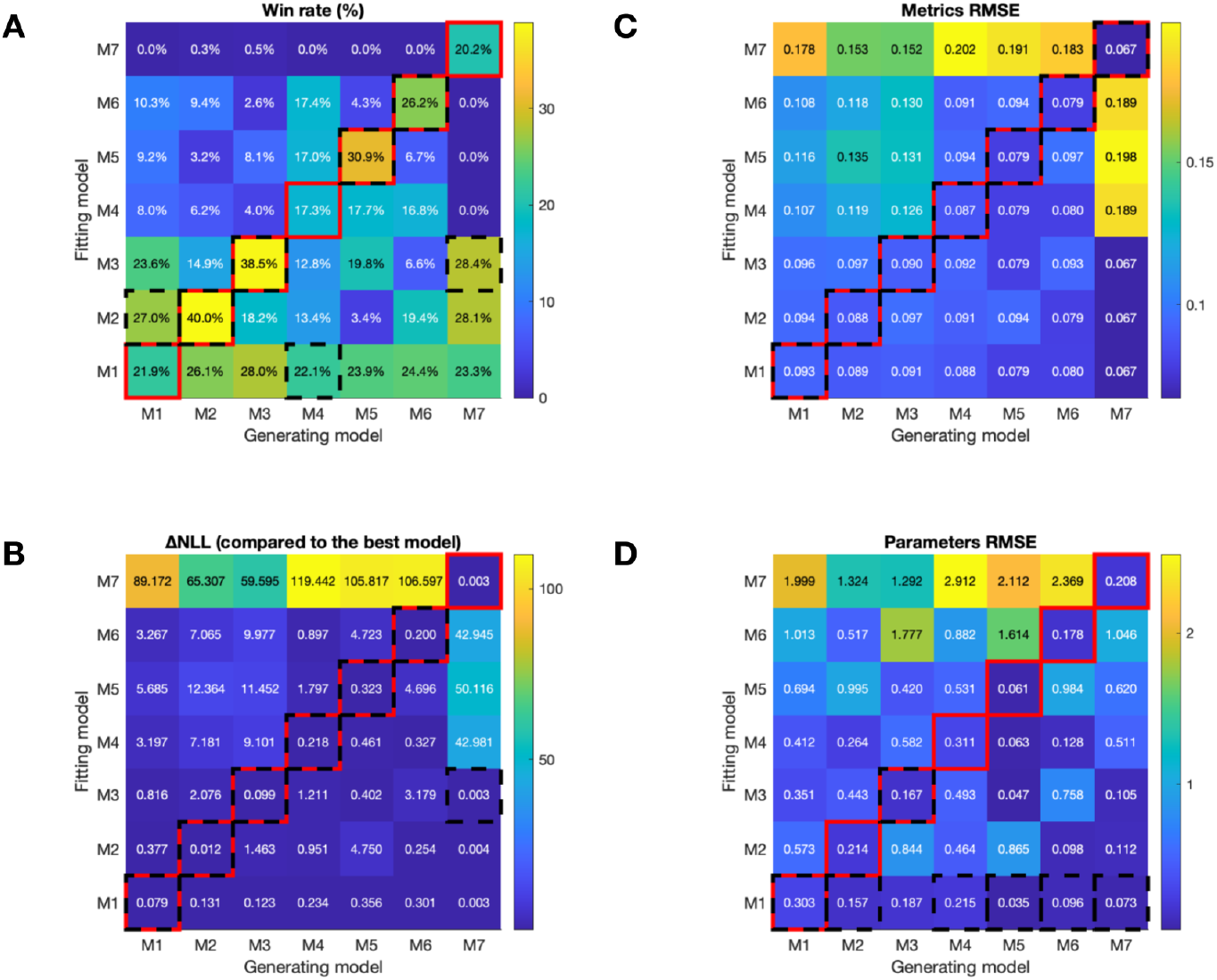
Model-recovery summary across generating and fitted models. Heatmaps use the same generating-model *×* fitted-model layout. Box notation follows the heatmaps in Figure S3. **(A)** Best-model win rate: percentage of iterations in which each fitted model achieved the lowest test negative log-likelihood (NLL), averaged across simulation conditions. **(B)** Difference in NLL from the best model: each model’s median test NLL minus the lowest median test NLL within the corresponding generating-model column. **(C)** Behavioral-metric RMSE: RMSE between simulated and predicted behavioral summaries, averaged across detection rate, accuracy, and response consistency. **(D)** Parameter RMSE: RMSE between ground-truth and recovered internal-noise parameters.

**Figure S6.**
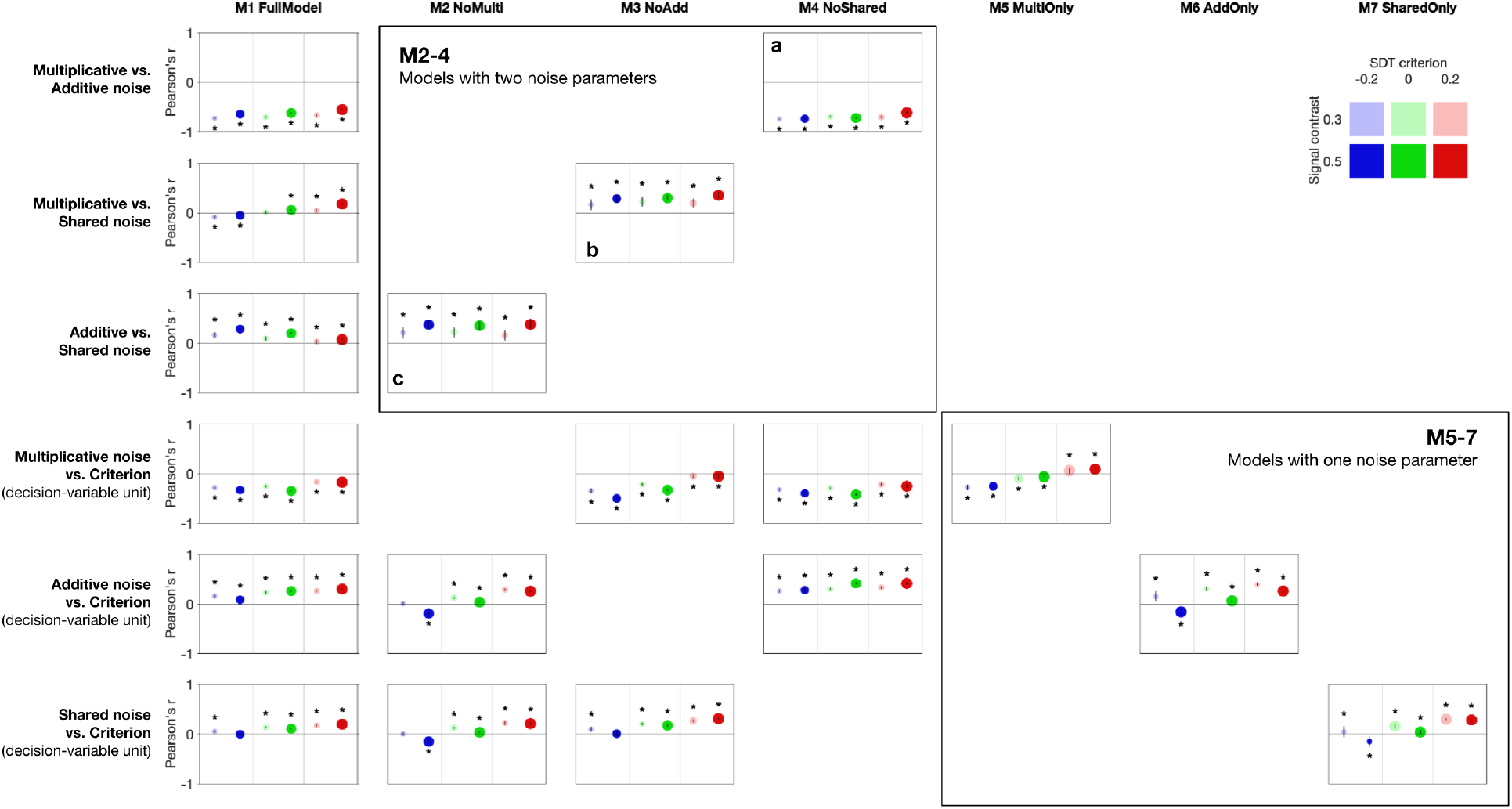
Parameter-pair correlations across matched-model simulations. Rows correspond to pairs of recovered parameters, and columns correspond to model variants. Only matched generating–fitted cases are included. Panels are omitted when one or both parameters were absent from the corresponding reduced model. In each panel, colored points show the Pearson correlation between two recovered parameters across fitting iterations for one signal-contrast *×* SDT-criterion condition, averaged across the included simulation conditions; error bars denote *±*1 SEM. Color encodes condition identity as in Figure S4. Vertical gray lines separate SDT-criterion groups, and point size is proportional to the number of simulation conditions contributing to each estimate. Asterisks indicate correlations outside the 95% permutation-null interval obtained from a two-tailed permutation test with 5,000 shuffles. The top three rows show correlations among internal-noise parameters, and the bottom three rows show correlations between each noise parameter and the decision criterion in decision-variable units. Black boxes highlight the correlations among pairs of internal-noise parameters.

## Acknowledgment

We thank Antoine Barbot, Aysun Duyar, Rania Ezzo, Marc Himmelberg, Yuna Kwak, Hsing-Hao Lee, and David Tu for their helpful comments.

## Author contributions

Conceptualization: MC, SX; Data curation: SX; Formal analysis: SX; Funding acquisition: MC, SX; Investigation: SX; Methodology: MC, MSL, SX; Project administration: MC; Software: SX; Supervision: MC, MSL; Visualization: SX; Writing–Original Draft Preparation: SX; Writing–Review and Editing: MC, MSL, SX.

## Data availability statement

The code used in this study is provided at https://github.com/shutianxue03/PF_RC. The processed data required to reproduce the figures are available at https://zenodo.org/uploads/21474441.

